# Mesenchymal-epithelial transition reduces proliferation but increases immune evasion in tumor spheroids

**DOI:** 10.1101/2024.09.23.614448

**Authors:** Gina Dimari, Yueyuan Hu, Annika Frenzel, Anke Fuchs, Alexander Wurm, Elisabeth Fischer-Friedrich

**Author notes:** Electronic mail: Corresponding author.

## Abstract

Mesenchymal-epithelial transition (MET) has been associated with secondary tumor outgrowth during metastasis but the underlying mechanism remains elusive. Using MET-inducible mesenchymal breast cancer cells, we investigated whether MET benefits tumor outgrowth by enhancing proliferation. We found that crowding inhibition of proliferation is present before and after MET, but mesenchymal cells gain a proliferative advantage through more effective escape from crowded cell islands. In 3D culture, proliferation is reduced upon MET with differential effects of focal-adhesion-signalling and actomyosin activity. In particular, inhibition of Src-signalling leads to increased growth after MET. Finally, in co-culture experiments, MET-induced tumor spheroids evade immune cell attack to a larger extent, likely due to more confined epithelial spheroid shape and changes in immunomodulatory molecules. Our data suggest that, contrary to previous assumptions in the field, MET might promote secondary tumor outgrowth not through a proliferation boost but through increased survival rate in the presence of immune cells.

## I. INTRODUCTION

EMT is a cellular transformation in which cells lose epithelial cell polarity and intercellular adhesion while gaining migratory potential^1^. Therefore, it has been linked to increased invasive capacity and the initiation of metastasis in cancer^2,3^. Nevertheless, studies show that secondary tumors arising at distant sites have increased epithelial features compared to EMT-transformed primary tumors, suggesting that the reverse transformation (MET) is required for successful initiation of metastatic growth^4–8^. However, the mechanisms by which MET promotes secondary tumor outgrowth remain elusive. There is evidence that the ability of the disseminated tumor cells to re-enter the cell cycle, proliferate and survive correlates with metastatic occurrence^9^. Therefore, one plausible mechanism is that MET increases cell proliferation. However, previous studies fail to confirm or disprove this concept. While some reports show that EMT leads to cell cycle arrest^10,11^ and that re-gaining E-cadherin expression enhances proliferation^12–14^; other studies describe that MET does not affect tumor initiating capacity or proves to be an adverse factor for tumor outgrowth^15–21^. Therefore, comprehensive studies examining the impact of MET on tumor cell proliferation under well-controlled conditions are crucial to unravel the complex factors influencing proliferation following MET.

Previous research shows that cell proliferation is strongly influenced by cell contacts with the extracellular matrix and junctions with neighboring cells^22–25^. The latter is of particular importance in epithelial cells where signaling through inter-cellular junctions restrains cellular proliferation through contact inhibition of proliferation^26^. The loss of contact inhibition is a hallmark of cancer progression and is tightly connected to the EMT process^27^. Focal adhesions are molecular complexes at the cell periphery that tether the actin cytoskeleton to the extracellular matrix via transmembrane integrin clusters. They are important regulators of proliferation and mechanotransduction through their connection to the actin cytoskeleton and signaling of focal-adhesion-associated kinases Src and FAK^28^. Focaladhesion-associated signaling is modified through cancerogenic transformations such as EMT and MET^28^. Previous studies show that Src and FAK can be stretch-activated through mechanical tension generated by actomyosin contractility, leading to a downstream increase of proliferative signaling^29^. Furthermore, FAK was shown to enhance the proliferation of endothelial cells through inhibition of the RhoA-ROCK pathway^30^.

Beyond cell proliferation, cell survival rate is a major factor that determines whether colonization is successful in the presence of mechanical and biochemical stresses as well as attacks from the immune system^31^. The EMT program has been reported to provide the cells with strategies to avoid immune system attacks mostly during intravasation and blood circulation^32^. However, this advantage of EMT-transformed mesenchymal cells may be lost at the micrometastasis stage. In fact, some studies show that EMT enhances anti-metastatic immune surveillance and makes cells more susceptible to the attack of natural killer cells^33,34^. In addition, higher levels of miR200c, a microRNA that inhibits EMT, has been associated to lower levels of tumor-infiltrating immune cells, specifically CD8+ T cells, and reduced overall survival of patients^35^. Furthermore, cell-cell junction protein expression levels are correlated with reduced infiltration of immune cells while high collagen cleavage by metalloproteinases (MMPs) is linked to increased immune cell invasion^36^. Accordingly, we hypothesize that differences in immune cell infiltration due to MET-related changes could be a decisive factor for the survival rate and growth of micrometastasis.

In this study, we investigated the effect of MET on proliferation using an inducible MET model of MDA-MB-231 breast cancer cells. These cells are known for their mesenchymal traits and their EMT signature^37–39^. In addition, MDA-MB-231 cells have proven to be highly metastatic when injected into mice^40–42^ what makes them a suitable candidate to model the outgrowth of a secondary tumor. In particular, we focused on cell junctions and focal-adhesions-mediated signaling using adherent and 3D cultures. Furthermore, we co-cultured tumor spheroids with peripheral blood mononuclear cells (PBMCs) to study the differences in immune cell infiltration and tumor cell survival. Interestingly, we found that both mesenchymal and MET-induced cells exhibit a mechanism of crowding inhibition that led to reduced proliferation at higher cell densities. However, mesenchymal cells gained a proliferative advantage through escaping crowded regions by migration. In 3D cultures, we found that MET caused an overall proliferation reduction in MDA-MB-231. In addition, we confirmed this key result in a second cell line - the mesenchymal ovarian cancer cell line ES-2. Further, we discovered that the role of focal-adhesion-associated kinases was changed by MET; mesenchymal cultures were particularly sensitive to inhibition of Src and actomyosin contractility, but MET-induced epithelial cultures were more affected by inhibition of FAK activity and its downstream target ROCK. Finally, we could show that while MET is a proliferative disadvantage in most culture conditions, the MET-induced epithelial state caused a reduction in immune infiltration of tumor spheroids leading to lower levels of apoptosis.

## II. RESULTS

### A. Overexpression of the miR-200c-141 microRNA cluster leads to MET in mesenchymal breast cancer cells

The miR-200 family of microRNA has been extensively linked to EMT in the context of cancer^43–46^. Previous studies showed that overexpression of two members from the family, miR-200c and miR-141, caused MET-characteristic changes in breast cancer cells^47,48^. Correspondingly, we generated a stable MDA-MB-231 miR-200c-141 overexpressing cell line by transfection with a doxycycline-inducible vector that codes for the miR-200c-141 cluster^47^. After induction, the cells displayed morphology and adhesion changes typical of MET, such as a cobblestone appearance and increased clustering (Fig. 1a). Further, qPCR confirmed the increase in miR-200c and miR-141 expression levels (Fig. 1b). Performing RNAseq analysis, we could show that not only the RNA of individual genes linked to an epithelial or mesenchymal phenotype changed their abundance accordingly (Fig. 1c, Supplementary Table 1), but also overall RNA changes show enrichment of genes that are typically downregulated by EMT according to GSEA analysis (Fig. 1d). Further, GSEA analysis shows a positive correlation of detected RNA changes with gene set enrichment of cell adhesion and cell junction proteins (Fig. S1g,h). In addition, we corroborated MET-induced changes of typical markers at the protein level with immunostainings and western-blotting: the epithelial marker E-cadherin was found to increase (Fig. 1e and S1a,b) while the mesenchymal transcription factor ZEB1 was downregulated upon miR-200c-141 overexpression (Fig. S1c,d,f). Vimentin and SNAI1 showed significant MET-induced changes of abundance on the RNA but not on the protein level (Fig. 1c and S1a-e).

**FIG. 1.**
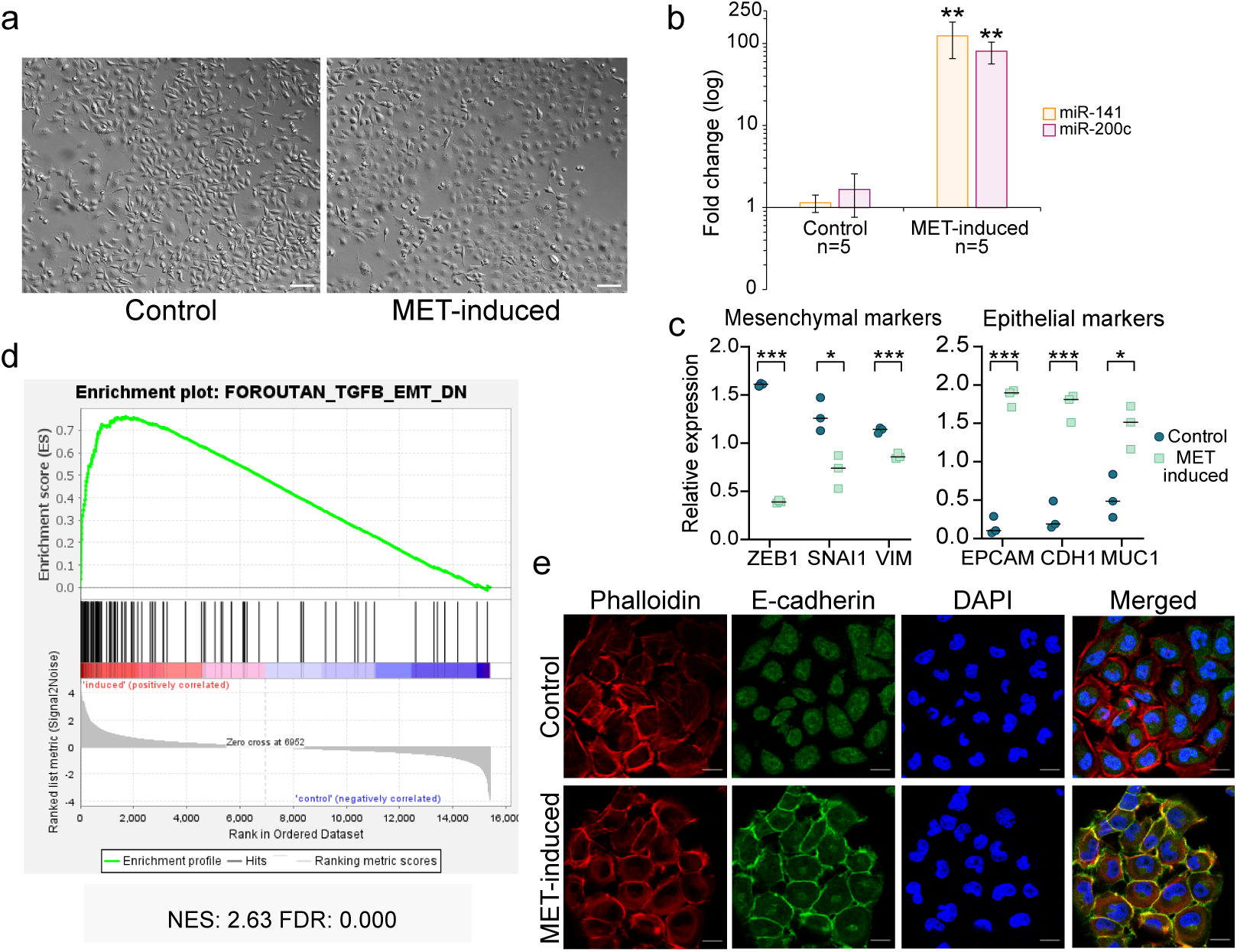
Overexpression of the miR-200c-141 microRNA cluster leads to MET in mesenchymal breast cancer cells MDA-MB-231. a) PlasDIC images of transfected MDA-MB-231 cells in 2D culture. Left: control cells without induction. Right: cells after 4 days of induction with 2 *µ*g/mL of doxycycline in the culture medium. Scale bars = 100 *µ*m. b) qPCR of relative expression fold changes upon induction shows a strong increase of microRNAs miR-141 and miR-200c. The number of biological replicates used for statistics is given by ‘n’. P-values for miR-141 and miR-200c fold changes were calculated using the Wilcoxon rank sum test comparing with the distribution of control fold changes. c) Changes in mRNA abundance of classical EMT markers upon induction quantified by RNA sequencing. Significance was tested with a two-tailed Student T-test and *p ≤* 0.05. d) GSEA enrichment plot of the human gene set “FOROUTAN TGFB EMT DN”^49^. NES = Normalized Enrichment Score, FDR = False Discovery Rate. e) Immunofluorescence of epithelial marker E-cadherin in uninduced cells (top panels) and cells after 4 days of induction (bottom panels). Scale bars = 20 *µ*m.

To examine the effects of MET on cell proliferation and survival, we utilize throughout this study MDA-MB-231 cells that contain the miR-200c-141 doxycycline-inducible vector unless specified otherwise.

### B. Increased cell density decreases proliferation in both mesenchymal and MET-induced MDA-MB-231 in 2D cultures

Given that cancer cells can grow as monolayers on surfaces and in three-dimensional clusters in the body^50^, we wanted to investigate proliferation differences upon MET in 2D and 3D culture. In the first place, we posed the question whether mesenchymal or MET-induced MDA-MB-231 breast cancer cells proliferate faster in 2D culture conditions. EMT-transformed mesenchymal cancer cells do not posses cell-cell junctions. Correspondingly, contact inhibition via cellular junctions and downstream Hippo signaling is lost as a mechanism of proliferation reduction when cells approach confluency^27,51,52^. We hypothesized that reverting EMT in mesenchymal MDA-MB-231 could restore contact inhibition by increased cell-cell adhesion leading to a corresponding proliferation decrease upon MET in high density cultures.

To test this hypothesis, we seeded control and MET-induced MDA-MB-231 cells in four defined densities. After three days of culture, we compared the effect of density on cellular proliferation and proliferative signaling through YAP1; nuclear accumulation of YAP1 promotes high proliferation and survival while cytoplasmic arrest of YAP1 is associated with cell growth inhibition^27,51^. The latter can be triggered e.g. via cell-junction-mediated contact inhibition^27,51^.

Therefore, we determined the ratio of YAP1 fluorescence in nuclei and cytoplasm; see Fig. 2a,b,e. Interestingly, in spite of a lack of classical adherens junctions proteins, mesenchymal MDA-MB-231 diminished YAP1 nuclear accumulation in response to a cell density increase in a similar manner as MET-induced epithelial cells (Fig. 2b,e). Only minor differences were detected at seeding density 1-3 with lower YAP1 nucleus/cytoplasm ratios in MET-induced cells (Fig. 2b).

**FIG. 2.**
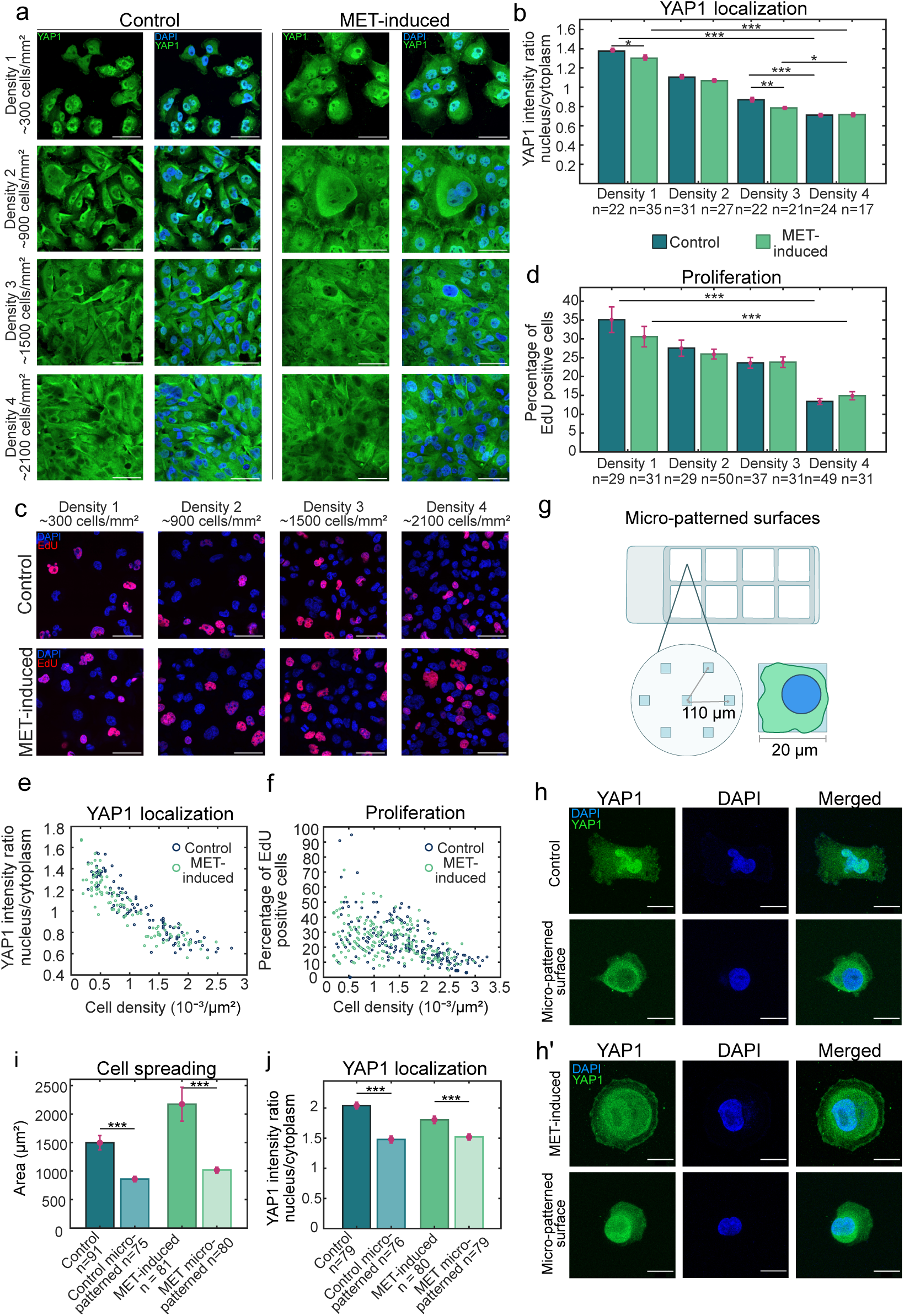
Cell crowding causes cytoplasmic arrest of YAP1 and reduced proliferation in both mesenchymal and MET-induced MDA-MB-231 in 2D culture. a) Immunofluorescence images showing the intracellular distribution of YAP1 (green) in control (mesenchymal) and induced (epithelial) cells at 4 different seeding densities after 3 days of growth. Nuclei are stained by DAPI (blue). Scale bar = 50 *µ*m. b) Nucleus-to-cytoplasm ratio of YAP1 intensity at different seeding densities corresponding to images shown in panel a. Average values of nucleus-to-cytoplasm ratios were calculated for each recorded frame, see panel a, giving rise to one data point in the statistics. For simplicity, the three-star significance between density 1 and 2, and density 2 and 3 were omitted in the chart. Details on the analysis can be found in the Materials and Methods Section IV L. c) Fluorescence images of cells subject to an EdU proliferation assay at 4 different seeding densities in control (mesenchymal) and induced (epithelial) cells. Cells that have undergone S phase during the incubation time show EdU positive nuclei (red) while the rest of the nuclei are stained with only DAPI (blue). Scale bar = 50 *µ*m. d) Percentage of EdU positive nuclei at different seeding densities for both conditions. For each recorded frame, fractions of EdU positive cells were determined. Correspondingly, each recorded frame gave rise to one data point, see Materials and Methods Section IV K 1. e, f) Scatter plots showing YAP1 localization ratio and percentage of EdU positive cells, respectively, in control and MET-induced MDA-MB-231 with relation to cell density. Cell density is expressed in number of nuclei per area of a recorded frame. g) Schematic representation of the micro-patterned surfaces used to constrain the area in which cells can adhere to the substrate on adhesive RGD squares of 20 *µ*m edge length. h, h’) Representative images of cells seeded on unpatterened (top row) and micro-patterned surfaces to limit the cell spread (bottom row). Top (h): control, bottom (h’): MET-induced. Green: YAP1 immunostaining, blue: DAPI staining of cell nuclei. Scale bar = 20 *µ*m. i, j) bar charts showing cellular spreading area (i) and YAP1 nuclear accumulation (j) for unconstrained cells and cells on micro-patterned surfaces. Each cell sampled constitutes a data point in the statistics. For panels b, d, i, and j, bar heights represent the mean and error bars show the corresponding standard error of the mean. Further, ‘n’ shows the total number of technical replicates sampled from at least 2 independent biological replicates. Significance was tested using a two-tailed Mann-Whitney-U-Test and *p≤* 0.05.

To also assess changes of cellular proliferation in response to increased cell density, we used an EdU incorporation assay, which stains cell nuclei that have undergone S-phase during the incubation time in red (Fig. 2c). By quantifying the percentage of EdU positive cells per frame, we were able to obtain a relative measurement of the proliferation rate for each seeding density (Fig. 2d,f). Similar to YAP1 nuclear accumulation, the percentage of EdU positive cells decreases as cell density increases for both mesenchymal and epithelial phenotypes (Fig. 2d,f).

We conclude that in spite of the absence of cell junctions, mesenchymal MDA-MB-231 cancer cells can still sense cellular crowding in cultures and reduce their proliferation accordingly.

### C. Reduction in spread area decreases nuclear YAP1 particularly in mesenchymal cells

Given that control MDA-MB-231 in their mesenchymal state showed a proliferation reduction upon crowding in the absence of canonical cell-cell junctions, we went on to test other factors that could influence YAP1 proliferative signaling. Increasing cell density corresponds to a reduced available cell spread area, which could in turn trigger a proliferation reduction. In fact, it has been previously shown that changes in the cell spread area can affect proliferation in certain cell types^23,53,54^. To test the influence of cell spread area, we constrained spread area by seeding cells in dishes with micro-patterned surfaces where cell adhesion is restricted to square patches coated with arginylglycylaspartic acid (RGD, a peptide that is present in many proteins of the ECM and which is recognized by cells as adhesion site^55^) of 20 *µ*m edge length (Fig. 2g,h,h’). As a control, cells were seeded with no spread area constraint at low densities on RGD-coated dishes (Fig. 2g,h,h’). With this approach, the effect of spread area reduction is disentangled from the increase in intercellular contacts in crowding-associated proliferative signaling.

We quantified the YAP1 nucleus/cytoplasm ratio after 24 h of culture time in conditions of constrained and unconstrained spread area. As expected, image analysis shows that spread area of cells cultured on micro-patterned surfaces was significantly lower with a more drastic reduction in the case of MET-induced cells (Fig. 2h,h’,i). Further, YAP1 nucleus/cytoplasm ratio declined significantly in correlation with reduced spread area in mesenchymal and epithelial cells, with a stronger decrease in mesenchymal cells (Fig. 2h,h’,j). Our findings suggest that reduced cell spread area inhibits proliferation in both mesenchymal and MET-induced epithelial MDA-MB-231 albeit with a more pronounced effect in mesenchymal MDA-MB-231. Therefore, cell crowding also inhibits proliferation in mesenchymal cells, at least in part, through a reduction in cell spread area at higher cell densities.

### D. Mesenchymal cells gain proliferative advantage through increased escape from crowded regions

EMT-transformed cells exhibit an enhanced migratory ability^1^. This change is most likely the main responsible factor for the influence of EMT on metastatic onset^2^. We asked how increased migration affects tumor cell proliferation in the presence of crowding inhibition of proliferation. To this end, we seeded cells in dense islands with open boundaries from which tumor cells can escape by de-adhesion and migration; see Fig. 3a,b. Within such islands, cells were more crowded in the center, than at the edges of the island for both islands of mesenchymal and MET-induced MDA-MB-231, see Fig. S3a,b,d and Fig. 3f.

**FIG. 3.**
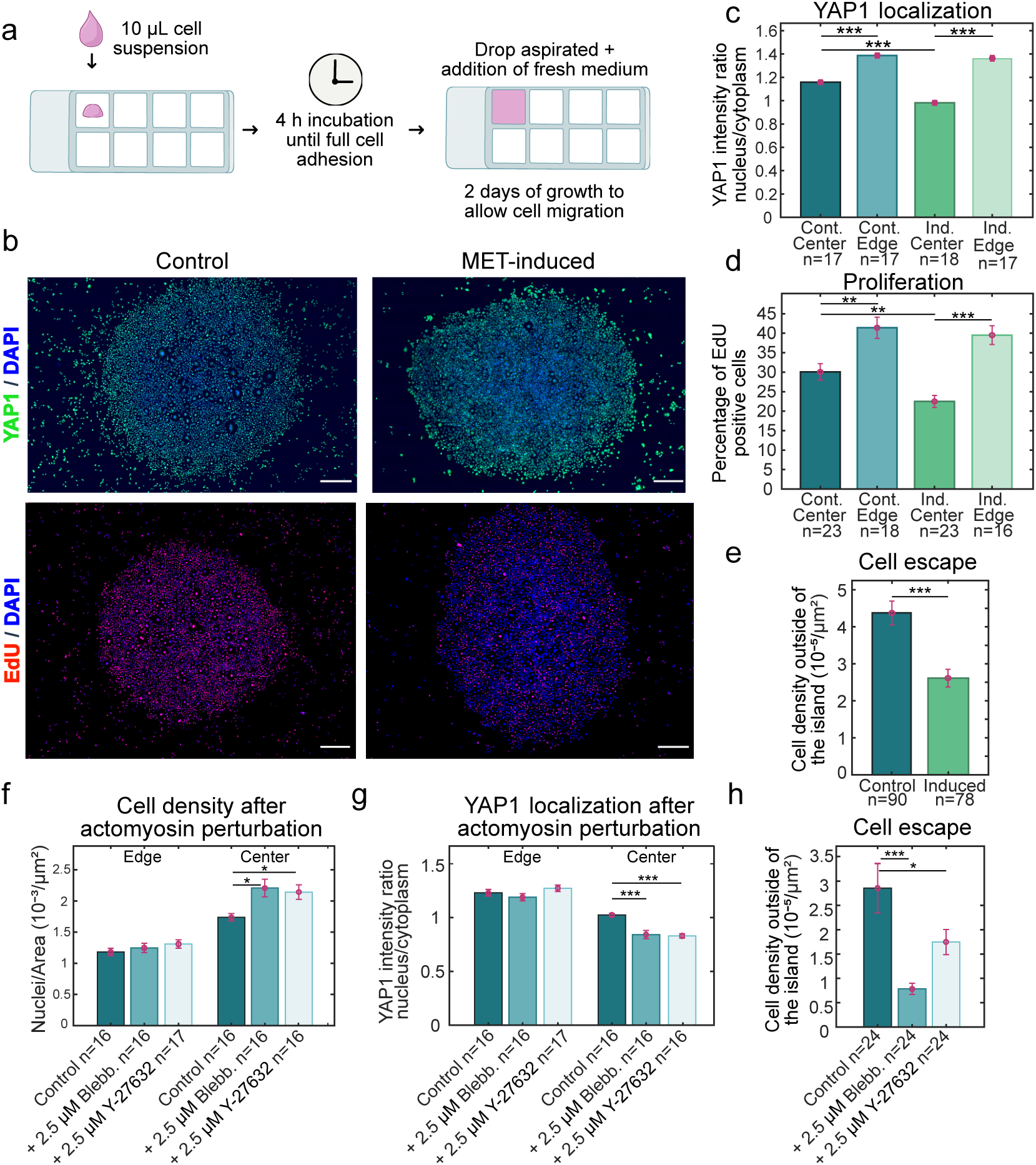
MET-induction reduces proliferative signaling in self-organized dense islands of MDA-MB-231 cells with open boundaries. a) Cells are seeded at high density in an isolated droplet of 10 *µ*L. After cell adhesion, additional medium is added to fill the well, see Materials and Methods Section IV M. b) Low magnification images of cell islands after two days of growth showing YAP1 localization (green, top row) and proliferation indicated by EdU-staining (red, bottom row). Cell nuclei are in addition stained with DAPI (blue), Scale bar = 500 *µ*m. c) YAP1 nucleus-to-cytoplasm ratio in center and border regions of adhered cell islands. d) Percentage of EdU-positive cell nuclei for both center and border regions. c,d) For each cell island, 4-5 frames were recorded for both center and edge regions. e) Cell nuclei density outside of cell islands. Each data point corresponds to cell counts in a peripheral region of on average 1 mm^2^ in a low magnification image, see Materials and Methods Section IV M. f,g,h) Actomyosin perturbation enhances crowding in the center of mesenchymal islands and reduces nuclear YAP1 and cell escape rates. Actomyosin was perturbed by the addition of: 2.5 *µ*M Blebbistatin or 2.5 *µ*M Y-27632 Rho Kinase inhibitor. For panels c, d, e, f, g and h, each data point corresponds to one recorded frame. The total number of recorded frames (technical replicates) is given by ‘n’. Data were obtained from 2 independent biological replicates that contained each 2 islands per condition. The height of the bars represent the mean for each condition and error bars show the corresponding standard error of the mean. Significance was tested using Mann-Whitney-U-Test and *p≤*0.05.

In a manner similar to our previous experiments, we assessed YAP1 subcellular distribution and cellular proliferation through immunostainings and an EdU assay, respectively. Fig. 3b shows low magnification confocal images of entire cell islands. However, for quantification purposes, higher magnification images were taken from areas defined as center or edge of an island (Fig. S3a,b). We found that for both phenotypes, control and MET-induced, YAP1 nuclear accumulation and proliferation were reduced in the center region compared to edge regions of the respective islands (Fig. 3c,d). Comparing the center regions between both phenotypes, we can appreciate that MET-induction leads to enhanced proliferative inhibition (Fig. 3c,d).

We hypothesized that mesenchymal MDA-MB-231 have an enhanced ability to migrate away from the crowded cell island and re-gain proliferative signaling. To confirm this, we quantified the number of cells in a peripheral range around the cell islands. We find a significantly higher number of cells outside cell islands of mesenchymal tumor cells (Fig. 3e).

To further test this hypothesis, we decided to hamper cell migration via inhibition of actomyosin activity using the myosin inhibitor Blebbistatin and the ROCK inhibitor Y-27632. After treatment with either of these inhibitors, we find on average higher densities with decreased nuclear YAP1 and proliferation of cells in center regions of the island when compared to untreated mesenchymal cells (Fig. 3f,g and S3f). In addition, this was accompanied by a lower number of cells that escaped from the islands (Fig. 3h). In contrast, analogous actomyosin inhibition applied to islands of MET-induced cells had no significant effects on cellular density or cellular escape from cell islands (Fig. S3d,e). However, nuclear YAP1 decreased in center regions (Fig. S3c). We speculate that this effect may be caused by the previously established dependence of the Hippo pathway on myosin II and ROCK activity^53^.

We conclude that mesenchymal cancer cells have a greater potential to escape cellular crowding through their enhanced migratory ability, thereby maintaining a proliferative state.

### E. MET reduces proliferation in MDA-MB-231 tumor spheroids

There is ample evidence that 3D platforms for cell culture provide a more physiological environment in both biochemical and biophysical terms for *in vitro* studies^56–58^. Therefore, we moved on to study MET-induced changes in proliferation in 3D-cultured tumor spheroids of MDA-MB-231 cells. To this end, we used PEG-Heparin hydrogels for which gel components are well-defined and for which stiffness and biodegradability can be adjusted^59^. This platform was previously established as a suitable scaffold for the growth of tumor spheroids from different cancer cell lines^20,60–62^.

First, we seeded cells within non-degradable PEG-Heparin hydrogels and cultured them for 14 days. Within this time, individual cells proliferated into multi-cellular tumor spheroids whose size and total cell number directly reflects the level of proliferation in control or MET-induced conditions. Contrary to what has been previously reported^4,12,14^, we find that MET induction leads to tumor spheroids with smaller cross-sectional areas, a reduced proliferation rate and a smaller number of cells with higher density after 14 days of growth (Fig. 4a-e).

**FIG. 4.**
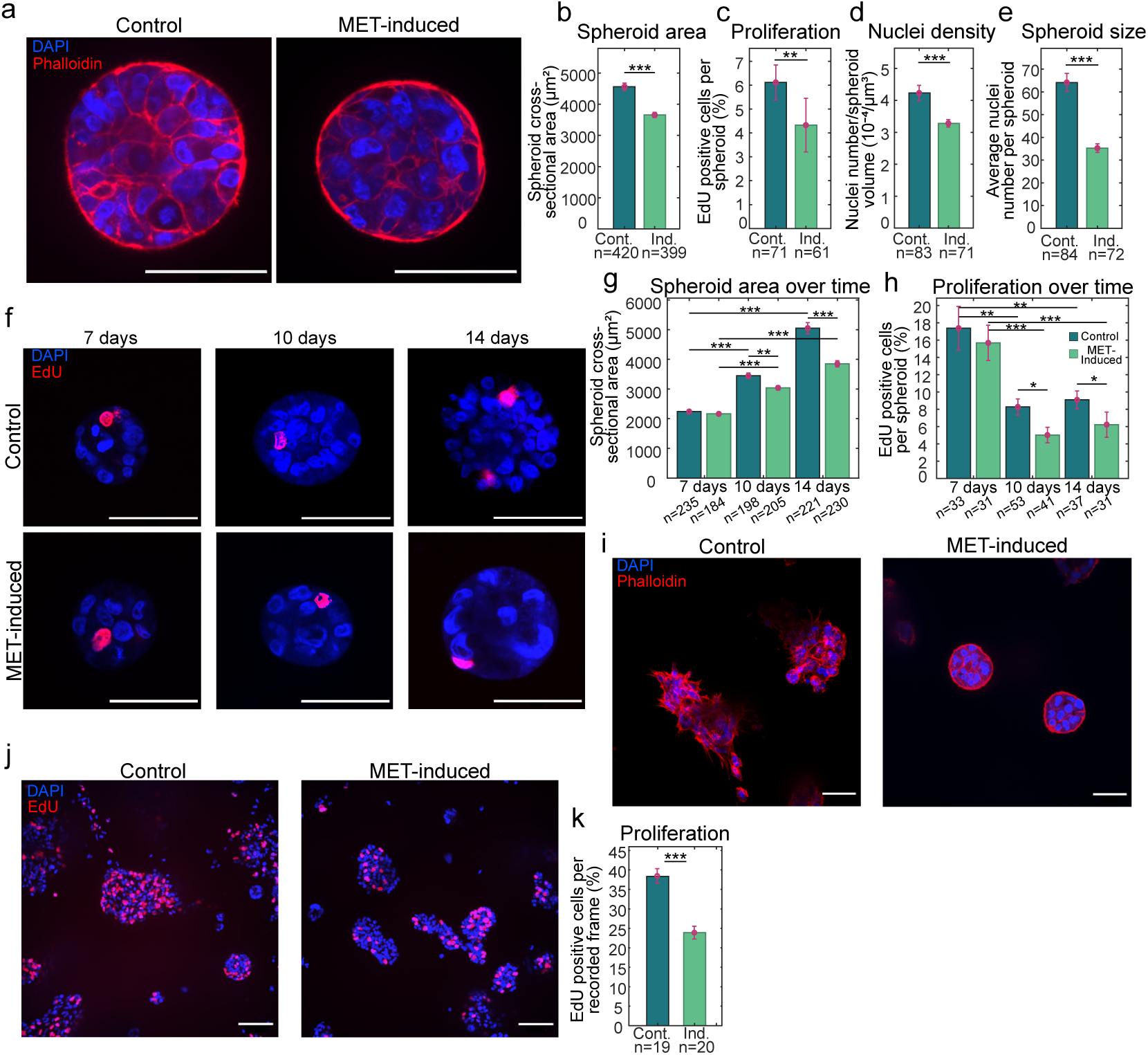
MET induction reduces proliferation of MDA-MB-231 tumor spheroids grown in PEG-Heparin hydrogels. a-h) Tumor spheroids cultured in non-degradable PEG-Heparin hydrogels. a) Representative fluorescence images of control and MET-induced tumor spheroid confocal cross-sections after 14 days of growth with DAPI staining for nuclei (blue) and phalloidin staining for f-actin (red). Scale bars = 50 *µ*m. b-e) Bar charts showing the quantification of b) maximal cross-sectional spheroid area, c) percentage of EdU positive cells, d) nuclei density and e) nuclei number per spheroid in 3D cultures of control and MET-induced MDA-MB-231 spheroids after 14 days of growth. Nuclei densities were calculated as the ratio between nuclei number and spheroid volume assuming sphere-shaped spheroids. Further, ‘n’ represents the total number of spheroids sampled from 4 independent biological replicates. Bar heights correspond to the mean and error bars show the standard error of the mean. Significance was tested using a two-tailed Mann-Whitney-U-Test and *p≤*0.05. f) Representative images of confocal cross-sections of spheroids subject to an EdU proliferation assay at 7, 10 and 14 days of growth for control and MET-induced cells. EdU-positive nuclei appear red and remaining nuclei are blue through DAPI-staining. Scale bars = 50 *µ*m. g,h) Maximal cross-sectional area (g) and percentage of EdU positive cells (h) quantified at 7, 10 and 14 days of growth for both control and induced MDA-MB-231 tumor spheroids. Data were obtained from 2 independent replicates. i-k) Tumor spheroids cultured in degradable PEG-Heparin hydrogels. i,j) Representative images of MDA-MB-231 cell clusters cultured for 10 days in control (left) and MET-induced (right) conditions. i) F-actin staining with phalloidin highlights morphological changes upon MET. Scale bars = 50 *µ*m. j) Cell clusters subject to an EdU proliferation assay. EdU-positive nuclei appear red and remaining nuclei are blue through DAPI-staining. Scale bars = 100 *µ*m. k) Percentage of EdU positive cells corresponding to conditions in panel j. The value of ‘n’ shows the total number of frames sampled from 2 independent biological replicates with at least 3 gels each. Bar heights correspond to the mean and error bars are the standard error of the mean. Significance was tested using a Mann-Whitney-U-Test and *p≤*0.05.

In addition, we monitored the dependence of spheroid growth rate on culture time. We find that after 7 days of culture, spheroid proliferation rates were similar in mesenchymal and MET-induced spheroids. For later culture times (10 and 14 days), proliferation rates decline in both conditions, likely due to increased cell crowding (Fig. 4f-h). This trend is stronger for MET-induced spheroids, demonstrating that MET enhances the progressive proliferation decline in confined tumor spheroids (Fig. 4h).

Furthermore, to mimic *in vivo* conditions more closely, we cultured MDA-MB-231 cells in degradable PEG-Heparin hydrogels. In these gels, PEG is modified so that it can be cleaved by MMPs, and thus the matrix can be remodelled by cells allowing for cell migration and unconfined growth^63^. While in control conditions mesenchymal MDA-MB-231 migrate through the gel forming loose star-shaped cell clusters (Fig. 4i, left), with MET-induction, the morphology of cell clusters is clearly distinct with approximately spherical shapes and no branches (Fig. 4i, right). Quantifying EdU positive cells, we find that MET induction also leads to a reduction of proliferation in degradable hydrogels (Fig. 4k). To verify that measured changes in cell proliferation stem from MET induction and not from possible off-target effects of the induction agent doxycyline, we confirmed that proliferation and spheroid size of untransfected MDA-MB-231 are not affected by the presence of doxycycline in PEG-Heparin gels (Fig. S4a-g).

We conclude that, opposite to common beliefs in the field, our results are consistent with MET leading to reduced tumor growth in both confined and unconfined 3D scaffolds in MDA-MB-231.

### F. Perturbation of focal-adhesion-associated kinases and actomyosin cytoskeleton affects proliferation differently on mesenchymal and MET-induced tumor spheroids

In the absence of RGD peptides in the hydrogels, we observed that, in both mesenchymal and MET-induced conditions, cells were barely able to proliferate. (Fig. S5a) Correspondingly, we concluded that integrin-associated signaling through focal adhesions plays a key role in spheroid proliferation. In addition, cell-ECM adhesion sites are known to be affected by the MET program^48^. Thus, we wanted to investigate the role of cell-ECM adhesion sites in proliferation differences between mesenchymal and MET-induced MDA-MB-231.

To perturb focal-adhesion signaling, we used pharmacological inhibitors against the two main kinases responsible for signal transduction: focal adhesion kinase (FAK) and the proto-oncogene tyrosine-protein kinase Src. Both kinases have been shown to be involved in proliferation of cancer cells^64,65^. We used FAK inhibitor 14 (also known as Y15) and Y11 to inhibit FAK^66^. Further, we employed Dasatinib and PP2 as inhibitors of Src kinase^67^. Inhibitors were added to the culture medium after one week of spheroid growth in non-degradable hydrogels and spheroids were harvested after one additional week of treatment. Measuring spheroid size and proliferation rate in terms of numbers of cell nuclei and EdU-labeling, respectively, we were able to identify changes in proliferation upon either FAK or Src inhibition; when inhibiting FAK in mesenchymal spheroids, only treatment with Y11 showed mild effects on mesenchymal tumor size and proliferation (Fig. 5a,b,e). In contrast, we observed a drastic effect when blocking Src activity with Dasatinib or PP2, leading to a strongly reduced spheroid size and proliferation (Fig. 5a,b,e).

**FIG. 5.**
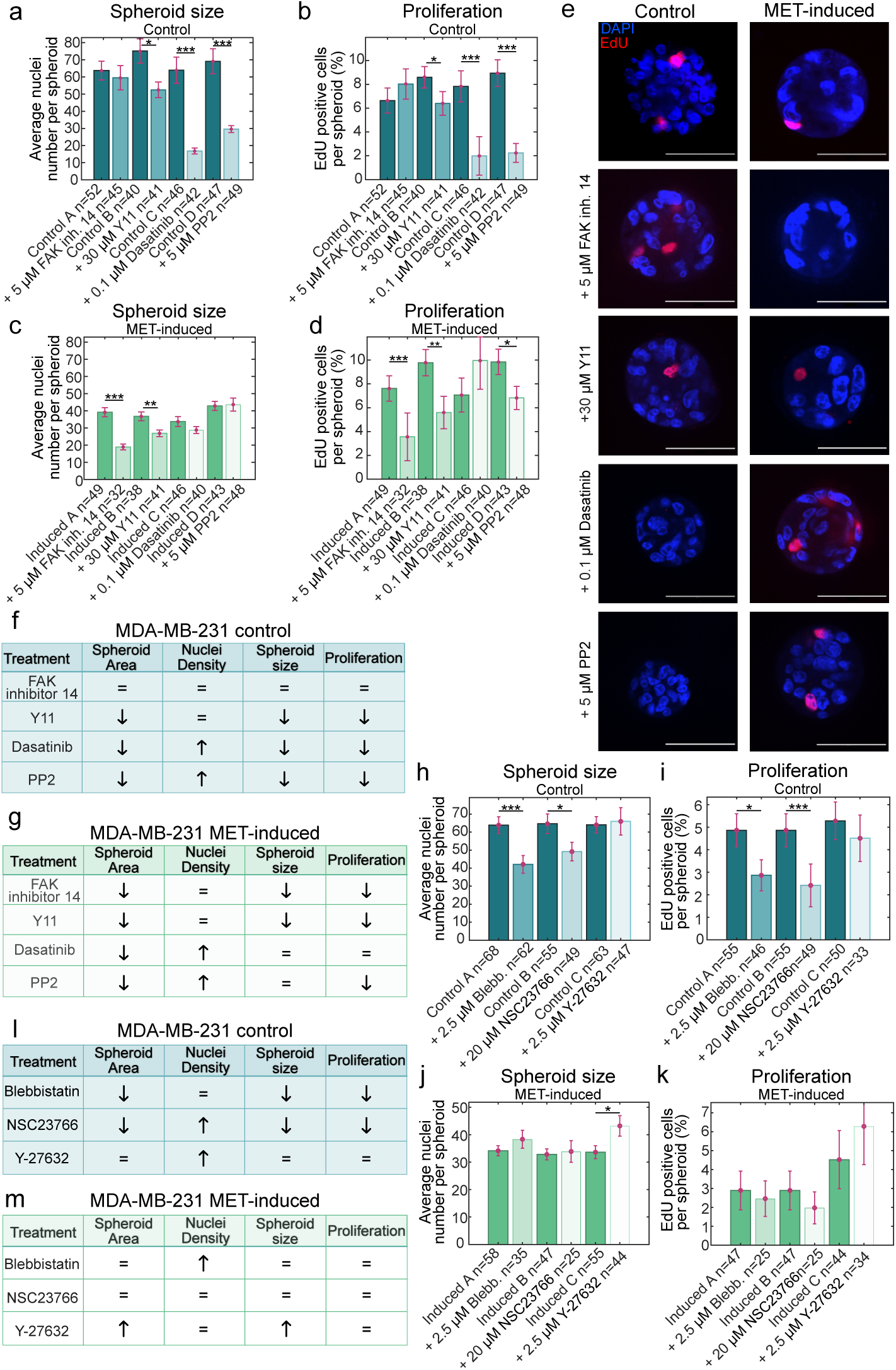
Inhibition of focal-adhesion-associated kinases and actomyosin affects proliferation differently in mesenchymal and MET-induced MDA-MB-231. a-d) Quantification of spheroid growth in terms of number of nuclei (a,c) and percentage of EdU positive cells (b,d) in tumor spheroids of control (a,b) and MET-induced (c,d) cells, upon inhibition of focal adhesion signaling molecules with: 5 *µ*M FAK inhibitor 14, 30 *µ*M Y11 (FAK inhibitor), 0.1 *µ*M Dasatinib (SRC inhibitor), and 5 *µ*M PP2 (SRC inhibitor).Control/Induced ‘A’, ‘B’, ‘C’ and ‘D’ are the control condition corresponding to each treatment. Bar heights correspond to the mean and error bars to the standard error of the mean. Significance was tested using a Mann-Whitney-U-Test and *p≤*0.05. The value of ‘n’ represents the total number of spheroids sampled; these were obtained from 2 independent biological replicates. e) Representative fluorescence images of tumor spheroids with and without signaling inhibition. EdU-positive nuclei are red and the rest of the nuclei in blue through DAPI-staining. Scale bars = 50 *µ*m. f,g) Summary of observed effects after inhibition of focal adhesion signaling molecules. (h-k) Spheroid size and proliferation quantified in terms of total nuclei number (h,j) and percentage of EdU positive cells (i,k) in tumor spheroids in control (h,i) and MET-induced (j,k) conditions. Where indicated, cytoskeletal perturbation of actomyosin contractility was applied with: 2.5 *µ*M Blebbistatin, 20 *µ*M NSC23766 RAC1 inhibitor, or 2.5 *µ*M Y-27632 Rho Kinase inhibitor. Control/Induced ‘A’, ‘B’, and ‘C’ are the control condition corresponding to each treatment. The value of ‘n’ represents the total number of spheroids sampled; these were obtained from 3 independent biological replicates. Bar heights correspond to the mean and error bars to the standard error of the mean. Significance was tested using the Mann-Whitney-U-Test and *p≤*0.05. l,m) Summary of observed effects after actomyosin inhibition.

In MET-induced MDA-MB-231 tumor spheroids, FAK inhibition with both inhibitors gave rise to a consistent decrease in spheroid size and proliferation rate (Fig. 5c,d,e). On the other hand, MET-induced MDA-MB-231 tumor spheroid size was unaffected by Src inhibitors Dasatinib and PP2 with a mild reduction in proliferation rate upon treatment with PP2 (Fig. 5c,d,e).

We also evaluated spheroid cross-sectional area and nuclei density with and without inhibition of focal-adhesion signaling, see Fig. S5b-e. Interestingly, we find that Src inhibition leads to reduction in spheroid cross-sectional area and an increase in nuclear density in mesenchymal and MET-induced spheroids. We suggest that this observation is connected to the role of Src in cell volume regulation^68^. A summary of all FAK- and Src-inhibition-induced spheroid changes is shown in Fig. 5f,g.

Focal adhesion complexes form the connection point of the actin cytoskeleton with the ECM^69^ and there is ample evidence for mechanotransduction and signaling cross-talk between both structures^70,71^. Given our findings that focal-adhesion signaling influences proliferation, we were interested in investigating if there are also changes in proliferation upon actomyosin inhibition. In particular, we chose to inhibit myosin II, RAC1 or ROCK activity, respectively, employing their inhibitors Blebbistatin, NSC23766 or Y-27632. Again, we grew spheroids in non-degradable hydrogels for one week and subsequently treated them with pharmacological inhibitors for one additional week. After this time, spheroids were fixed, stained and imaged to analyze changes in proliferation (Fig. S5j).

Interestingly, the results show that inhibiting myosin II and RAC1 interfere with the proliferation of mesenchymal spheroids, but have no effect on MET-induced spheroids (Fig. 5h-k). Only ROCK inhibition via Y-27632 treatment showed an effect on MET-induced spheroids, surprisingly leading to an increase in spheroid size and cross-sectional area (Fig. 5j and S5g). We note that an analogous effect upon ROCK inhibition has been reported previously for spheroids of epithelial breast cancer cells^62^. This effect might be accounted for by the interaction between ROCK and the cell cycle regulator PAK1 (also p21)^62,72^. In Fig. 5l,m, we display a summary of the changes in spheroid parameters measured upon actomyosin inhibition.

In conclusion, our results suggest that MET affects the nature of cell proliferation regulation; while mesenchymal cells rely largely on Src kinase activity and actomyosin integrity to promote growth, MET-induced spheroids rely more on FAK and ROCK activity to maintain proliferation. We speculate that these actomyosin-mediated proliferation changes are connected to our observations of Src- and FAK-mediated proliferation changes; stretch-activation of Src by actomyosin-mediated tension and FAK-mediated inhibition of the RhoA-ROCK pathway were shown to be connected to proliferative signaling^29,30,73^.

Finally, we note that Src inhibition can lead to a situation where MET-induced tumor spheroids gain proliferative advantage over mesenchymal tumor spheroids.

### G. MET reduces apoptosis rate of tumor spheroids in the presence of peripheral blood mononuclear cells (PBMCs)

Tumor growth is not only influenced by cell proliferation rates but also by cell survival probabilities^9^. Immune surveillance presents a major obstacle to cancer cell survival in the colonization phase^31^. Thus, we went on to test the increase of apoptosis in tumor spheroids in the presence of immune cells. We hypothesized that the morphological differences between tumor cell clusters formed by MDA-MB-231 cells before and after MET could lead to differences in immune cell infiltration and consequently the survival rate of the tumor cells. In particular, mesenchymal cell clusters were star-shaped with a large surface-to-volume ratio, while MET-induced epithelial cell clusters formed dense confined spheroids in degradable hydrogels, see Fig. 4i.

To test this hypothesis, we co-cultured MDA-MB-231 tumor spheroids with PBMCs obtained from a donor blood sample. This subpopulation of immune cells consist of lymphocytes (T cells, B cells, and Natural killer cells), monocytes, and dendritic cells. The T cell population comprises both CD4+ and CD8+ T cells^74^. This composition is a suitable immune cell population to induce an antitumor immune response which is carried out mainly by CD8+ T cells and Natural killer cells aided by antigen presenting cells^75^.

Following previously established approaches^76,77^, we grew tumor clusters from MDA-MB-231 in matrigel, both in control and MET-inducing conditions for 10 days followed by a 3-day co-culture with PBMCs. After this time, apoptosis and infiltration of PBMCs were measured. To quantify apoptotic levels in breast cancer cells after co-culture, we used confocal imaging of fixed and stained co-culture samples as well as flow cytometric analysis of dissociated live stained cells from co-cultures. Fig. 6a shows representative confocal images of the morphology in matrigel 3D-cultures with colocalized PBMCs (red dots). To identify apoptotic cells and PBMC infiltration in confocal images, an antibody against Cleaved Caspase-3 and DAPI nuclear staining was used while PBMCs were identified by their prior labeling with cell proliferation dye eFluor 670 (Fig. 6b). In this way, we quantified the number of apoptotic foci and infiltrating PBMCs per tumor cell cluster, see Fig. 6c,d, S6e,f. The results show that higher levels of apoptosis were found upon the addition of PBMCs in the cultures for both conditions. However this increase was significantly higher in mesenchymal MDA-MB-231 tumor clusters (Fig. 6c). Concomitantly, we find that a larger number of PBMCs had infiltrated cancer cell clusters (Fig. 6d).

**FIG. 6.**
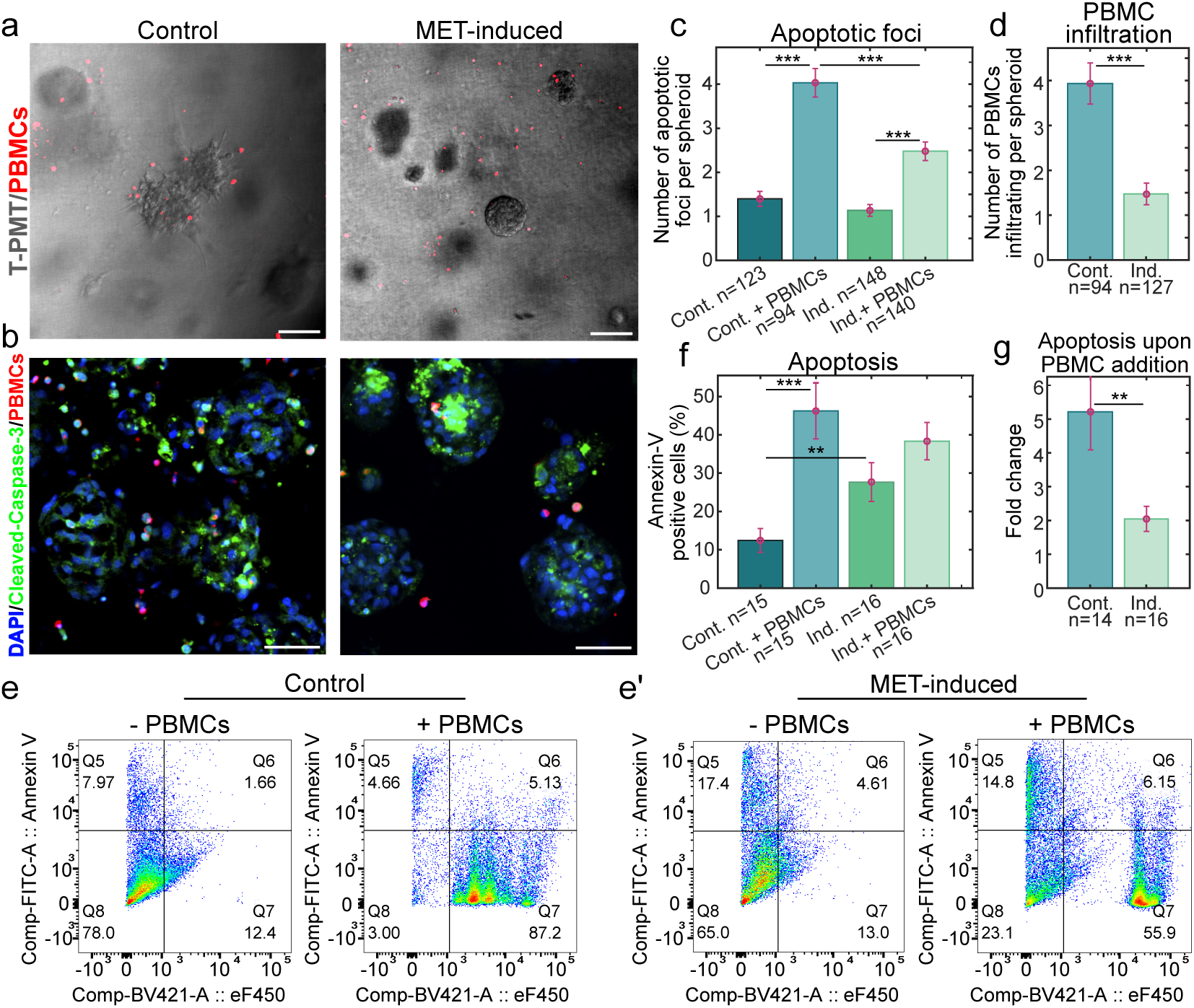
MET enhances survival rate of tumor spheroids in the presence of peripheral blood mononuclear cells. a) Confocal images of transmitted light and red fluorescence of MDA-MB-231 tumor spheroids in control (left) and MET-induced (right) conditions after 10 days of growth and 3 days of co-culture with PBMCs. PBMCs appear as red dots due to labeling with the cell proliferation dye eFluor 670. T-PMT = transmitted light detection module. Scale bar = 100 *µ*m. b) Representative confocal images of fixed tumor spheroids in co-culture with PBMCs (blue: DAPI-stained cell nuclei, green: immunofluorescence of Cleaved Caspase-3 as apoptotic marker, red: eFluor 670-stained PBMCs). Scale bar = 50 *µ*m. c, d) Quantification of apoptotic foci and spheroid-infiltrating PBMCs from confocal images of fixed and stained spheroids such as shown in panel b (See Materials and Methods section IV O). The value of ‘n’ represents the total number of clusters sampled; these were obtained from 3 independent biological replicates. e) Representative plots from flow cytometric detection of MDA-MB-231 cells in coculture with PBMCs. Propidium Iodide-staining was used to detect the non-necrotic cell population, fluorescence intensity of cell proliferation dye eFluor 670 or 450 (x-axis) was used to identify PBMCs, fluorescence intensity of FITC-Annexin-V (y-axis) to detect apoptotic cells in the sample. The percentage of apoptotic tumor cells were calculated using the ratio of values from quadrant 5 and 8, see panel f. f) Percentage of Annexin-V positive cells in the tumor cell population of the co-cultures from the flow cytometry analysis. The value of ‘n’ represents the total number of technical replicates; these were obtained from 3 independent biological replicates. g) Fold change increase in apoptosis upon the addition of the PBMCs to the culture. The wells with and without PBMCs were paired according to their position in the well plate to obtain a fold change. For c), d), f) and g), bars correspond to the mean and error bars are standard error of the mean. Significance was tested using the Mann-Whitney-U-Test and *p≤*0.05.

For flow cytometry measurements, the matrigel was degraded and the tumor spheroids were disaggregated into single cells. During flow cytometric analysis, we first used propidium iodide (PI) to detect viable cells, Annexin-V staining to mark apoptotic cells and eFluor 450/670 fluorescence to identify PBMCs (Fig. S6a,a’ and Fig. 6e). In this way, we were able to gate for the apoptotic fraction of cancer cells. Quantification of the percentages of apoptotic cells in mesenchymal and MET-induced tumor spheroids are shown in Fig. 6 f,g. Similar to the results obtained from confocal imaging of fixed samples, we observed significantly higher apoptosis levels upon PBMC addition in mesenchymal control MDA-MB-231 (Fig. 6f,g).

These results suggest that the metastatic success of MET-induced cells could be at least partially explained by an improved immune evasion. This evasion is promoted by the more compact, spatially confined shape of the cell clusters and reduced cluster surface area after MET which leads to a lower encounter rate with PBMC.

In addition, we find that upon MET-induction MDA-MB-231 have increased mRNA levels of the immune checkpoint programmed death ligand-1 (PD-L1) (Fig. S6g), a mechanism better known to suppress T cell responses^78^. This was observed in conjuction with the downregulation of the mRNA of three clinically relevant immune co-stimulatory molecules: CD40, TNFSF18 (also known as GITRL), and TNFSF9 (also known as 4-1BBL) (Fig. S6i) which in turn are known to activate T cells and lead to anti-tumor immunity^79–81^. Therefore, an MET-induced change in expression of immunomodulatory proteins may additionally contribute to greater immune evasion after MET in MDA-MB-231 cells.

### H. MET reduces proliferation of ovarian cancer tumor spheroids

Our previous experiments have given evidence of a decrease in cell proliferation upon MET induction in mesenchymal breast cancer cells MDA-MB-231. Since this finding challenges the idea of MET being required for outgrowth of secondary tumors, we wanted to test if it can be reproduced in a second biological model system. In ovarian cancer, the incidence of EMT has been extensively reported and linked to chemotherapy resistance, tumor progression, and stemness^82,83^. In addition, MET in ovarian cancer has been linked to macro-metastasis formation^84^. Correspondingly, we decided to investigate if inducing MET in post-EMT ovarian cancer cells would lead to a similar proliferation reduction as in MDA-MB-231 cells. For this, we chose the mesenchymal ovarian cancer cell line ES-2. Through a stable transfection of this cell line with the doxycycline-inducible miR-200c-141 pINDUCER13 vector, we could overexpress microRNAs miR-141 and miR-200c causing clear genetic and morphological changes in line with MET both in adherent cultures and 3D cultures in degradable hydrogels (Fig. 7a-e, Fig. S7a-c, Supplementary Table 2). We used an EdU incorporation assay to measure proliferation changes in degradable hydrogel cultures analogous to our assay in MDA-MB-231 described in Fig. 4i,j. Quantification shows that MET-induction leads to a reduction in proliferation in unconfined ovarian cancer tumor spheroids (Fig. 7f,g). As a control, we verified that doxycycline addition had no effect on proliferation of untransfected ES-2 cells (Fig. S7d,e). We conclude that our experiments with ovarian ES-2 cells confirm an MET-induced proliferation decline in the same manner as in breast cancer cells MDA-MB-231.

**FIG. 7.**
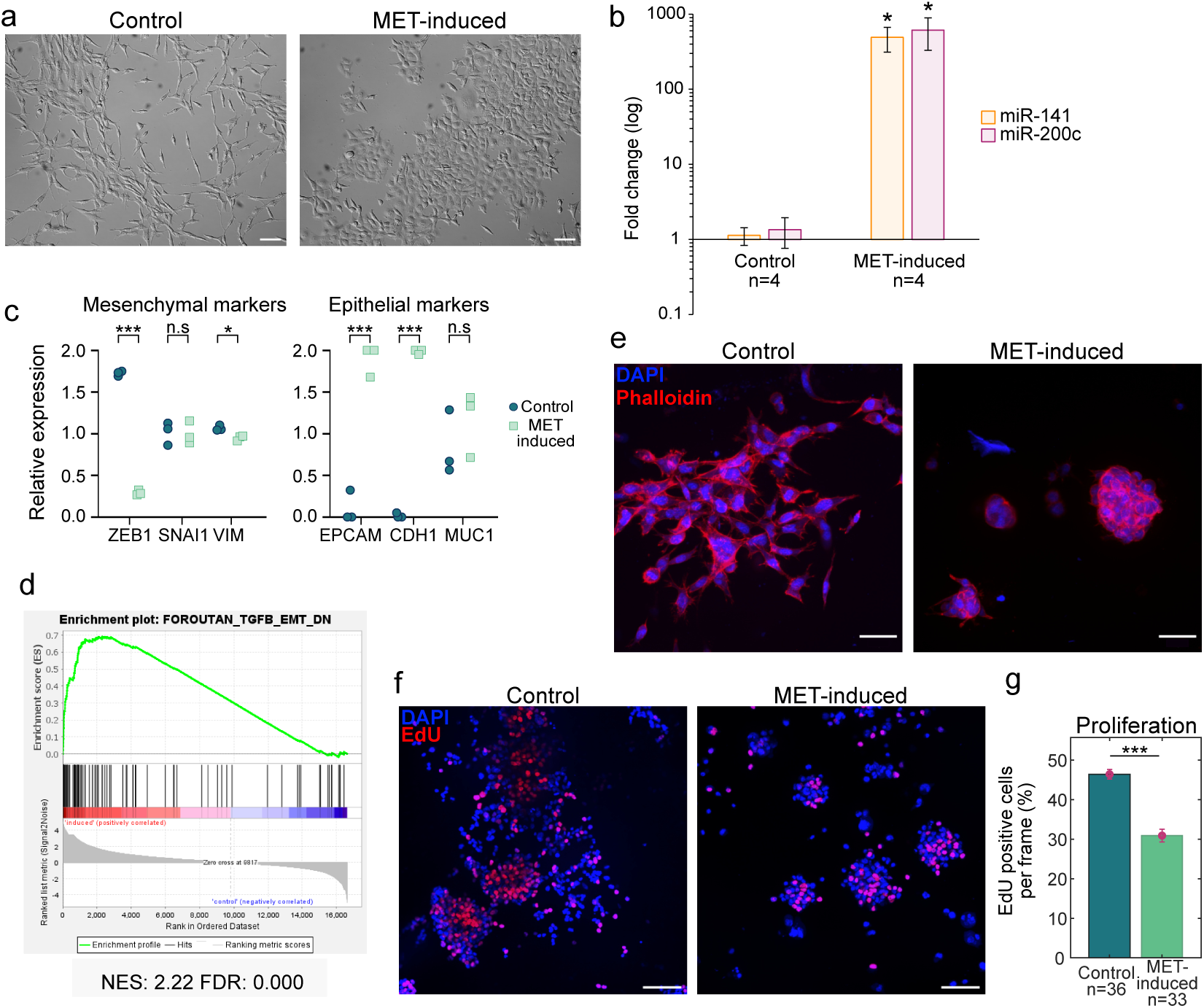
Overexpression of miR-200c-141 induces MET and reduces proliferation in mesenchymal ovarian cancer cells ES-2. a) PlasDIC images of transfected ES-2 cells in adherent cultures. Left: control cells without induction. Right: cells after 5 days of induction with 2 *µ*g/mL of doxycycline in the culture medium. Scale bar = 100 *µ*m. b) qPCR of microRNAs miR-141 and miR-200c shows a strong increase upon induction. P-values were calculated using the Wilcoxon rank sum test. Here, ‘n’ represents the total number of biological replicates. c) Changes in mRNA abundance of classical EMT markers upon induction quantified by RNA sequencing. Significance was tested with a two-tailed Student T-test and *p ≤* 0.05. d) GSEA enrichment plot of the human gene set “FOROUTAN TGFB EMT DN”^49^. e) Representative images of morphological changes observed upon MET in degradable PEG-Heparin hydrogels. Left: control, right: MET-induced (right) cells grown in degradable hydrogels for 10 days, fixed and stained with phalloidin (red) and DAPI (blue). Scale bar = 50 *µ*m. f) Representative confocal images of control (left) and MET-induced (right) ES-2 cells grown in degradable hydrogels for 10 days and stained with the proliferation marker EdU (red) and DAPI (nuclei, blue). Scale bar = 100 *µ*m. g) Percentage of EdU positive cells corresponding to conditions presented in panel e. Here, ‘n’ represents the total number of frames sampled; these were obtained from 2 independent biological replicates. Bar heights correspond to the mean and error bars are standard error of the mean. Significance was tested using a Mann-Whitney-U-Test and *p≤*0.05.

## III. DISCUSSION

Previous research suggests that MET is a key factor for the outgrowth of secondary tumors during the metastatic colonization of new tissues. To elucidate the mechanisms by which MET influences secondary tumor outgrowth, we investigated how MET affects cellular proliferation and immune evasion in the mesenchymal breast cancer cell line MDA-MB-231.

In the first part of our study, we explored the role of proliferation inhibition through cell crowding in adherent cultures in mesenchymal and MET-induced conditions. Interestingly, we find that cell crowding caused a similar reduction in proliferation in both control and MET-induced cells. While this is not surprising for MET-induced epithelial cells in which E-cadherin and cell-cell contacts are re-established^52^, see Fig. 1c,e and S1a, b, the underlying mechanism is less clear for mesenchymal conditions. Our data suggest that mesenchymal, and also to a lesser extent MET-induced cells, experience a decline in proliferative signaling in response to reduction in spread area, featuring cell-substrate contact as an essential proliferative factor. As reduced spread areas naturally emerge at high cell densities, we propose that this mechanism is a potent means of proliferation inhibition in mesenchymal MDA-MB-231 upon cellular crowding. While we find that proliferation rates for mesenchymal and MET-induced cells are on a similar level at a certain cell density, see Fig. 2, we observed that mesenchymal cells gain proliferative advantage through escape from crowded regions, see Fig. 3. Blocking actomyosin contractility in mesenchymal cells, we find impeded cell escape and concomitantly reduced proliferation while MET-induced cells were unaffected by this treatment, see Fig. 3.

By growing cells in the more physiological conditions of 3D cultures, we saw an overall proliferation reduction caused by MET. This key finding was corroborated in a second mesenchymal cell line of ovarian cancer, see Fig. 7. We went on to study the influence of cell-matrix adhesion on proliferation and identified focal-adhesion-associated kinase Src and actomyosin contractility as major regulators of the growth of mesenchymal tumor spheroids. In contrast, the growth of MET-induced tumor spheroids was more strongly affected by Src-independent FAK activity and its downstream effector ROCK. In conclusion, our data on 3D cultures show that cell proliferation depends on signaling mediated by focal adhesions for both mesenchymal and MET-induced cells, although the concrete signaling pathways are different in the two cases.

Taken together, our results portray an overall proliferation reduction after MET. This conclusion is supported by the findings of some earlier studies. Kim et al. reported that E-cadherin expression in mesenchymal MDA-MB-231 leads to Hippo pathway mediated contact-inhibition and corresponding proliferation reduction in crowded cultures^52^. Injecting MDA-MB-231 cells into mice, Olmeda et al.^18^ showed that MET induced through SNAI1-silencing led to reduced tumor incidence and growth rate and that SNAI1 expression was required for lymph node metastasis. In addition, Ran et al.^19^ induced MET in MDA-MB-231 by depleting CBF*β* and reported reduced tumor growth at the site of injection and lower number of metastatic events when cells were injected into the circulation. Correspondingly, we conclude that there is increasing evidence that MET may be an adverse factor to proliferation.

In the final part of our study, we addressed the effect of immune cell interaction with tumor cell cultures in mesenchymal and MET-induced conditions. Geometrical considerations predict that in the absence of chemotactic signaling, immune cell encounter of a cancer cell cluster is proportional to the cluster surface area. As MET-induction significantly reduces cluster surface area through increased cell adhesion and spatial cluster confinement, see Fig. 6a and Fig. 4i, a declined encounter rate is expected. In accordance with this consideration, we find in our study that in the presence of PBMCs, tumor cell infiltration is more abundant in mesenchymal cancer cell clusters. Concomitantly, apoptotic rates are more substantially increased than in MET-induced cultures, see Fig. 6. The observed MET-induced evasion of immune cell attack in our assay may be in part mediated through expression changes of immunomodulatory proteins; we observed in the transcriptome the downregulation of immune co-stimulatory molecules upon MET and further confirmed the upregulation of the immunosuppressing molecule PD-L1, see Fig. S6. We note that in our assay an alloreactive response of immune cells was triggered due to genetic incompatibility of the PBMC donor and MDA-MB-231 breast cancer cells. In conclusion, our findings suggest that the role of MET in metastatic success could be explained by enhanced survival of immune attacks at the point of colonization of distant tissues at the stage where secondary tumors are still small. Our findings are in contrast to previous studies that reported enhanced immune escape of tumor cells and increased expression of PD-L1 upon EMT^32,85^.

In conclusion, our study challenges the common idea that MET facilitates secondary tumor outgrowth during metastasis through an increase of cell proliferation. Instead, our data supports the alternative hypothesis that MET increases cell survival in the presence of immune cells through a more compact shape of micro-metastasis and changes in the expression of immunomodulatory proteins.

## IV. MATERIALS AND METHODS

### A. Cell culture

MDA-MB-231 cells stably transfected with the construct pINDUCER13 miR-200c-141 were cultured on DMEM with 10% Fetal Bovine Serum (#10270106, Thermofisher), 1% Pen/Strep (#15140122, Thermofisher) and 0.5 *µ*g/mL Puromycin at 37 °C with 5% CO_2_. ES-2 cells stably transfected with the construct pINDUCER13 miR-200c-141 were cultured on DMEM with 10% Fetal Bovine Serum (#10270106, Thermofisher), 1% Pen/Strep (#15140122, Thermofisher) and 1 *µ*g/mL Puromycin at 37 °C with 5% CO_2_. For both cell lines, medium was exchanged every 2-3 days and cells were split when 70-90% confluency was reached. To induce MET transformation in cells, doxycycline (Sigma Aldrich #D3072) was added to the medium at a final concentration of 2 *µ*g/mL during at least 4 days. In this way, the expression of microRNA cluster miR-200c-141 was induced.

### B. Plasmids and transfection

MDA-MB-231 cells were stably transfected with pINDUCER13 miR-200c-141 using Turbofectin (OrgiGene #TF81001) according to manufacturer’s protocol. ES-2 cells were stably transduced with pINDUCER13 miR-200c-141 by lentiviral transduction. For Lentivirus production, packaging (psPAX2), envelope (pMD2G.VSV.G), and corresponding transfer plasmids (pINDUCER13 miR-200c-141) were transfected into HEK293T cells using PEIpro transfection reagent (Polyplus, Illkirch, France) according to the manufacturer’s protocol. Harvested lentiviral particles and 8 µg/ml polybrene were used for transduction of ES-2 cells. Selection was performed using 3 µg/ml Puromycin for 96 hours, followed by a constant treatment with 1 µg/ml Puromycin. The plasmid was a gift from Jeffrey Rosen & Thomas Westbrook (Addgene plasmid #81020; http://n2t.net/addgene:81020; RRID:Addgene 81020).

### C. Immunostaining for EMT markers and transcription factors

Adherent cells were fixed with 3.7% formaldehyde/PBS for 10 minutes, followed by a 10 minutes permeabilization step with 0.2% Triton X-100 (#T8787, Sigma Aldrich). The cells were then blocked for 1 hour at room temperature with 5% Goat Serum/PBS. The cells were then treated with primary antibody of Vimentin (#5741T, Cell Signaling) at a dilution of 1:100, E-cadherin (#60335, ProteinTech) at a dilution of 1:50, ZEB1 (#21544-1-AP, Proteintech) at a dilution of 1:100, YAP1 (#13584-1-AP, ProteinTech) at a dilution of 1:100, SNAI1 (#26183-1-AP, ProteinTech) at a dilution of 1:50 overnight at 4 *^◦^*C in 1% BSA/PBS. Cells were then treated with the corresponding secondary Alexa Fluor 488 highly cross adsorbed conjugate (#A32731, #A32723, Invitrogen) at a concentration of 1:2000 in 2% BSA/PBS for 1.5 h at room temperature. Afterwards, the cells were treated with 5 *µ*g/mL DAPI (#28718-90-3, Sigma Aldrich) and 0.2 *µ*g/mL phalloidin-iFluor-647 (#ab176759, Abcam) for 10 minutes in 2% BSA/PBS solution. In between every step, the sample was washed three times with PBS. Images were taken with either a Zeiss LSM700 confocal microscope with a 40x/1.2 C-Apochromat water immersion Zeiss objective or an ND1 Nikkon spinning disk confocal microscope with a 40x/1.15 Apo LWD water immersion Nikon objective both from the Physics of Life microscopy facility.

### D. 3D cultures

#### 1. Non-degradable PEG-Heparin hydrogels

As described before^61,62^, in situ assembling biohybrid multiarmed poly(ethylene glycol) (starPEG)-heparin hydrogels, prepared as previously reported^86^, were used for embedding cells to grow into spheroids. In brief, cells were detached from the tissue flasks and resuspended in PBS. Subsequently, they were mixed with a freshly prepared heparin-maleimide-8 solution to obtain a final cell density of *≈* 10^6^ cells per mL (to reach a final concentration of 0.5 *×* 10^6^ cells/mL in the gel) with a heparin-maleimide-8 concentration of 3 mM. RGD (Arginylglycylaspartic acid) was added in a concentration of 3.5 *µ*g/*µ*L to the heparin-cell solution. For a molar ratio, PEG/heparin-maleimide of gamma = 0.5, PEG precursors (MW 16000) were reconstituted at a concentration of 1.5 mM and put in an ultrasonic bath for 10–20 s in order to dissolve. Then, 20 *µ*L of the heparin-cell suspension was mixed with equal volume of PEG solution in a pre-chilled Eppendorf tube, using a low binding pipette tip. Thereafter, a 20 *µ*L drop of PEG-Heparin-cell mixture was pipetted onto an ibidi 8-well slide suitable for imaging (#80826, ibidi) and cured for a gelation period of around 4 minutes before adding the medium. Final concentrations of PEG and HM8 were correspondingly 1.5 mM and 0.75 mM. Using AFM indentation of hydrogels with cantilevers (CSC37/tipless/no aluminum, Mikromasch) modified with polystyrene beads of 10 *µ*m diameter, emergent Young’s moduli of these gel were determined to be *≈* 1000 Pa^62^.

The gels were cultured for 14 days with 300 *µ*L of medium that was exchanged every other day. For wells that were supposed to give rise to MET-induced spheroids, doxycycline was supplemented to the medium at a concentration of 2 *µ*g/mL.

#### 2. Degradable PEG-Heparin hydrogels

Similar to non-degradable gels (see above), cells were detached from the tissue flasks and resuspended in PBS and mixed with the prepared heparin-maleimide-8 solution of the same concentration as for the non-degradable gels, also at a density of *≈* 10^6^ cells per mL. PEG precursors were reconstituted at a concentration of 2.2 mM (for a molar ratio PEG/heparin-maleimide of gamma=0.75) with a PBS at a pH 6.5 and put in an ultrasonic bath for 10–20 s in order to dissolve. 20 *µ*L of the heparin cell suspension were mixed with equal volume of PEG solution in a pre-chilled Eppendorf tube, using a low binding pipette tip. Thereafter, a 20 *µ*L drop of PEG-heparin-cell mixture was pipetted onto an ibidi 8-well slide suitable for imaging (#80826, ibidi) and waited for a gelation period of 8 minutes before adding the medium. Final concentrations of PEG and HM8 were correspondingly 1.5 mM and 1.1 mM. The gels were cultured for 10 days with 300 *µ*L of medium that was exchanged every other day.

#### 3. Matrigel

For spheroid cultures in matrigel, Geltrex (#A1413201, ThermoFischer) was mixed with the culture medium at a 20% volume fraction. Both the medium and the matrigel along with the eppendorf tubes used were kept on ice for the whole mixing procedure. 6000 cells were re-suspended with a total of 100 *µ*L of the cold Geltrex-medium solution and seeded in 96 well plates with optical bottom (#655090, Greiner bio one). The wells were briefly incubated on ice after the seeding of the cells for *≈*10 min so that most of the cells would settle at the bottom of the dish and grow closer to the glass slide facilitating later imaging. The spheroids were cultured for approximately 10 days and medium was exchanged every 2 days.

#### 4. Fixation and imaging of spheroids/clusters in hydrogels

For imaging, the PEG-heparin hydrogel cultures were fixed with 3.7% formaldehyde/PBS for 1 h, permeabilized with 0.2% Triton X-100 for 30 minutes and stained with 5 µg/mL DAPI (#28718-90-3, Sigma Aldrich) and 0.2 µg/mL phalloidin-iFluor-647 (#ab176759, Abcam) for 1 hour in 2% BSA/PBS solution. Imaging was done with either a Zeiss LSM700 confocal microscope with a 40x/1.2 C-Apochromat water immersion Zeiss objective or a ND1 Nikkon spinning disk confocal microscope with a 40x/1.15 Apo LWD water immersion Nikon objective both from the PoL Light Microscopy Facility.

### E. Western blotting

Protein expression differences upon MET were analyzed using western blots, see Fig. S1. Cells were seeded onto a 6-well plate and grown up to a confluency of 80–90% with or without MET-inducing agents. Thereafter, cells were lysed in SDS sample/lysis buffer (62.5 mM TrisHCl pH 6.8, 2% SDS, 10% Glycerol, 50 mM DTT and 0.01% Bromophenol blue). The lysates were then incubated at 4 *^◦^*C for 30 minutes in a rocking shaker followed by a short sonication step of 2 minutes and a denaturation at 95 *^◦^*C for 10 minutes. 20 µL of the cleared lysate were then used for immunoblotting. The cleared lysates were first run on precast protein gels 4-20% or 7.5% (#456-1096 or 456-1093, Bio-Rad) in MOPS SDS running buffer (B0001, Invitrogen). Subsequently, proteins were transferred to nitrocellulose membranes (GE10600012, Sigma-Aldrich). Nitrocellulose membranes were blocked with 5% (w/v) skimmed milk powder (T145.1, Carl Roth, Karlsruhe, Germany) in TBST (20 mM Tris–HCl, 137 mM NaCl, 0.1% Tween 20 (pH 7.6)) for 1 h at room temperature followed by washing with TBST, and incubation at 4 *^◦^*C overnight with the corresponding primary antibody diluted 1:1000 E-cadherin (#60335, Proteintech), 1:10.000 Vimentin (#60330, Proteintech), 1:1000 ZEB1 (#21544-1-AP, Proteintech), 1:2000 YAP1 (#13584-1-AP, ProteinTech), 1:500 SNAI1 (#26183-1-AP, ProteinTech) and 1:5000 GAPDH (#ab9485, Abcam) in in 5% (w/v) bovine serum albumin/TBST solution. Afterwards, the blots were incubated with appropriate secondary antibodies conjugated to horseradish peroxidase, Goat anti-mouse HRP (#ab97023, Abcam) or Goat anti-rabbit HRP (#ab97051, Abcam) at 1:5000 dilution in 5% (w/v) skimmed milk powder in TBST for 1 h at room temperature. After TBST washings, specifically bound antibodies were detected using Pierce enhanced chemiluminescence substrate (ECL) (PN: 32109, Invitrogen). The bands were visualized and analyzed using an iBright FL1500 Imaging System (#A44241, ThermoFisher). For statistics, three individual biological replicates were used. The intensity values were normalized towards the housekeeping gene values, in this case GAPDH.

### F. RNA extraction

RNA purification for real-time PCR and RNA sequencing was done from cell pellets containing (at least) 0.5 *×* 10^6^ cells using the RNeasy Plus kit (#74134, Qiagen) following the manufacturer’s protocol.

### G. microRNA detection by quantitative real-time PCR

Both the cDNA synthesis and the quantitative PCR reactions were performed with the products from the Custom TaqMan*^T^ ^M^* Small RNA Assay kit (#4398988, ThermoFischer) according to the manufacturer’s protocol. All the primers were obtained also within the TaqMan assay for the specified microRNA. RNU6B was used for normalization. The details of the master mixes prepared and the cycling steps for each reaction can be found in Tables S3-S6. The quantitative reaction was run with a BioRad CFX96 real time cycler. For statistics, three individual biological replicates were used and three technical replicates were obtained in each experiment. The analysis of the relative expression was done by the ΔΔCT method.

### H. RNA sequencing

#### 1. Library preparation for RNA sequencing

RNA sequencing (RNA-seq) libraries were prepared using the Stranded mRNA Prep, Ligation Kit (Illumina) according to the manufacturer’s instructions. Briefly, mRNA was purified from 1 µg total RNA using oligo(dT) beads, poly(A)+ RNA was fragmented to 150 bp and converted into cDNA, and cDNA fragments were end-repaired, adenylated on the 3’ end, adapter-ligated, and amplified with 12 cycles of PCR. The final libraries were validated using a Qubit 2.0 Fluorometer (Life Technologies) and a Bioanalyzer 2100 system. All barcoded libraries were pooled and sequenced 2×75bp paired-end on an Illumina NextSeq550 platform to obtain a minimum of 10 Mio reads per sample.

#### 2. Standard workflow for RNA sequencing data analysis

In the context of the RNA-seq data analysis workflow, quality control assessment of raw reads was performed using *FastQC*, followed by *Trimmomatic* trimming to remove low-quality sequences. The trimmed reads were then aligned to the reference genome using *STAR version 2.7.9a*. To facilitate the mapping process, a genome reference index was constructed using either *GRCh37.fa (hg19)* or *GRCh38.108.gtf (hg38)* as the reference. During the alignment step, specific parameters were employed to ensure accurate and reliable alignment results. These parameters included: alignIntronMax (maximum intron size) set to 500000; alignMatesGapMax (maximum gap between paired mates) 500000; outFilterMultimapNmax (maximum number of multiple alignments allowed) 100; outFilterMismatchNoverLmax (maximum ratio of mismatches allowed) 0.05; chimSegmentMin (minimum length of chimeric segment) 15; chimScoreMin (minimum chimeric score) 1; chimJunctionOverhangMin (minimum overhang for chimeric junctions) 15; chimSegmentReadGapMax (maximum gap in reads within a chimeric segment) 3; and alignSJstitchMismatchNmax (maximum mismatch in stitched junctions) set to 5 -1 5 5. The output was converted to sorted BAM files with *SAMtools*. Differentially expressed genes (DEG) were calculated using the ratio of the mean expression of normalized read counts (RPKM - Reads per kilobase of transcript per Million mapped reads) from induced versus non-induced cells for all annotated genes. These normalized values were then used for gene set enrichment analysis and plotted as log_2_ fold change (LFC) against the negative log_10_ of the p-value within a 2D-plot using pandas.

### I. Gene set enrichment analysis

Gene expression data were analyzed for enrichment using GSEA software (Broad institute version 4.3.2)^87,88^ and MSigDB (version MSigDB v2022.1.Hs, September 2022)^89,90^. Gene expression data were loaded into GSEA in the format specified by the software user guide. The permutation type chosen was “Gene Set” as directed in the GSEA user guide when having fewer than 7 samples per phenotype, and permutation number was set at 1000 for testing significance. The dataset was collapsed to symbols before the analysis. The chip platform used was ‘Human Gene Symbol with Remapping MSigDB.v2023.1.Hs.chip’. A p-value and FDR of less than 0.05 was considered significant.

### J. Coating of glassware

#### 1. Fibronectin coatings

Glass bottom slides used for YAP1 localization experiments and EdU stainings (see Fig. 2, 3, S2 and S3) were coated with fibronectin. The surface of each well was covered with a solution of Fibronectin (#F1141, Sigma-Aldrich) of 20 *µ*g/mL in PBS and incubated at 37 *^◦^*C for 1 h. Afterwards, the remaining solution was aspirated and the surface was left uncovered in a sterile environment to air dry for approximately 30 minutes. Seedings were done immediately afterwards.

#### 2. RGD coatings

The surface of each well was covered with a solution of RGD of 10 *µ*g/mL in PBS and incubated at 37°C for 1 h. Afterwards, the remaining solution was aspirated and the surface was washed with sterile distilled water and left uncovered in a sterile environment to air dry for approximately 30 minutes. Seedings were done immediately afterwards.

### K. Quantifications in 2D and 3D proliferation assays

#### 1. EdU incorporation assays

EdU (#BCK647-IV-IM-S, Sigma-Aldrich) at a final concentration of 5 *µ*M was added to the cultures according to the manufacturer’s protocol and incubated for, either 3 h for adherent cultures or 4 h for 3D cultures in hydrogels prior to fixation. Following fixation, permeabilization and EdU click reaction were carried according to the manufacturer’s protocol. In addition, DAPI staining was done to the samples as described before.

#### 2. Nuclei quantification for 2D and 3D EdU proliferation assays

To quantify nuclei numbers in both 2D and 3D samples the Image J ‘trackmate’ plug-in was used^91^. Beforehand, images were processed by binning to a pixel size of *≈* 1 *µ*m and by applying a Gaussian blur filter. The DoG detector was used to detect the objects in the stacks and the conditions. Software parameters were adjusted to detect observed nuclei in different experiments setting ‘object diameter’ between 12-15 *µ*m and ‘quality threshold’ between 10-50.

To quantify spheroid nuclei density, the nuclei numbers were divided by the volume of the spheroid (calculated from the cross-sectional area from the fluorescence image, assuming a sphere-shaped spheroid).

#### 3. Quantification of tumor spheroid cross-sectional area

Cross-sectional areas of tumor spheroids in non-degradable hydrogels were quantified with Fiji using transmitted light images of spheroids obtained from a Zeiss Axio Vert.A1 microscope using a A-Plan 10x/0.25 Ph1 Zeiss objective.

### L. YAP1 localization in dependence of cell densities in 2D culture

#### 1. Cell seeding at different densities

Induced cells and untreated cells were seeded into different densities (60, 300, 600 and 1500 cells*/*mm^2^) in fibronectin-coated 18 well glass bottom slides (#81817, ibidi) and kept in culture for 3 days with daily medium change. On the third day, cultures were fixed and immuno-stained.

#### 2. Fluorescence intensity analysis

For the YAP1 quantification as presented in Fig. 2, a customized ImageJ macro was used (https://gitlab.com/ polffgroup/met analysis). In essence, the analysis uses nuclear and cytoplasmic masks to measure the average immunofluorescence of YAP1 in adherent cells in the nuclear areas and in the cytoplasmic regions (without nucleus) in the image. Then, the ratio between the two intensities is calculated. To that end, a mask of the nuclear region was obtained first from thresholding the images using the DAPI channel. Then, a mask for cellular regions was obtained by thresholding the YAP1. In the next step, a cytoplasmic mask was obtained by subtracting the nuclear mask from the cellular mask. The output of the ImageJ macro are the integrated fluorescence intensities of Yap1 immunofluorescence *I*_nuc_ and *I*_cyt_ in the nuclear or cytoplasmic regions, respectively. Furthermore, the macro outputs the values 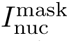 and 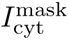 corresponding to the intensity of the nuclear and cytoplasmic masks which corresponds to the respective positive area multiplied by 255. Then, the average fluorescence intensity of Yap1 in the nuclear and cytoplasmic regions were obtained by normalizing the immunofluorescence values towards the positive area of each mask, i.e. as 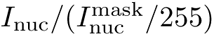 and 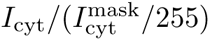.

For Fig. 2, we also determined the number of nuclei in each recorded confocal image (185 *×* 185 *µ*m^2^, 1152 *×* 1152 pixel^2^) to calculate the local nuclear density of the chosen field of view in the culture. For this purpose, we used the ImageJ plugin ‘Analyse particles’. Nuclear densities were grouped according to the following intvervals of number of nuclei per frame: 0-20, 20-40, 40-60, 60-100. The interval centers correspond approximately to nuclear densities as specified in Fig. 2a-d,, i.e. 300, 900, 1500, 2100 cells*/*mm^2^. In panels Fig. 2c,d, each recorded frame gave rise to one data point in the statistics that gave rise to the plotted bar charts.

### M. Self-assembled cell islands

For the seeding, cell suspensions of both control and induced cells in a concentration of 2*×*10^6^ cells/mL were prepared. 10 *µ*L drops of cell suspension were placed in the center of a well of a fibronectin-coated 8-well ibidi glass bottom slide (#80827, ibidi). The drops were incubated for 4 hours for full cell attachment to take place. Afterwards, the medium from the drop was aspirated and the well was washed once with fresh medium to get rid of unattached cells. Then, 300 *µ*L of the respective fresh medium were added to the well. Subsequently, the cells were cultured for two days. Fixation and staining were performed as described above. For the analysis of the cell island experiments, confocal images were taken from center and edge regions of the islands, see Supplementary Fig. S3. The analysis of the YAP1 signal ratio and the EdU percentage was done as described above.

Large field images of the whole island were recorded by using a 10x objective (Nikon Plan Apo 10x/0.45 DIC N1) and stitching several independent images together was done automatically by the — ‘Large image’ option of the Nikon software, see Fig. 3. For the quantification of the cells in the periphery, six different approximately equidistant regions in the environment of an island were selected with areas between 0.5 *−* 2 mm^2^. Regions were chosen such that they had a minimum of 200 *µ*m distance from the outer island edge and a maximum of 800 *µ*m. Half of the regions were chosen on the left side of the island, while the remaining half was located on the right-hand side of the island. The area of each region was measured along with the number of nuclei in the region. Nuclei were counted by thresholding of the DAPI channel of the image and subsequent application of the ‘Analyze particles’ plugin of ImageJ.

### N. Area constraint experiments

For constraining the area in which the cells can spread and make contact with the substrate, the cells were seeded at low densities on RGD micro-patterned surfaces squares with a side length of 20 *µ*m (#83801-S, ibidi). For comparison, cells were seeded at the same low density on normal RGD-coated glass slides (#80827, ibidi) that did not constrict spreading (see Sec. IV J 2). For seeding, cells in either condition were detached and re-suspended in a concentration ranging from 1.7 *×* 10^4^ to 3.3 *×* 10^4^ per mL. Then, 300 *µ*L of cell suspension was added to each well of the respective slide to achieve a density between 5 *×* 10^3^ and 10 *×* 10^3^ cells per well. After 23 h of culture, EdU was incorporated in the culture media (see Sec. IV K 1) for 1 h and afterwards the cells were fixed and stained as described in previous sections.

### O. Co-culture of MDA-MB-231 tumor spheroids with Peripheral Blood Mononuclear cells (PBMCs)

#### 1. Peripheral Blood Mononuclear Cell isolation

Buffy coat preparations were obtained as by-product from red blood cell donations from the German Red Cross blood donation center in Dresden (DRK Nord-Ost gGmbH) after written consent of donors. Peripheral Blood Mononuclear Cells (PBMCs) were isolated from buffy coat by the standard density gradient isolation. Briefly, 50 mL Leucosep tubes were loaded with 15 mL Lymphoprep™ (STEMCELL Technologies), centrifuged at 1000 g for 1 minute at room temperature (RT) to let Lymphoprep™ pass through the porous barrier. The blood was 1:1 diluted with sterile PBS and 30 mL of diluted blood were carefully overlaid into the Leucosep tube using a serological pipette and centrifuged at 800 g for 40 minutes at RT. After centrifuging, the PBMCs (opaque interphase) were carefully collected and transferred into a new 50 mL falcon tube using a Pasteur pipette, followed by washing with PBS at 500 g for 10 minutes at RT with a subsequent wash with PBS at 200 g for 10 minutes at RT. The cell pellet was re-suspended in PBS for cell counting. The PBMCs were then pelleted and resuspended in CryoStor® CS10 freezing medium (#07930, STEMCELL Technologies). The freezing cell suspension were aliquoted into cryovials and then placed into CoolCell® Containers and kept at -80 °C for more than 1 day. Thereafter the cryovials were transferred into liquid nitrogen tanks.

#### 2. Co-culture

MDA-MB-231 tumor spheroids were cultured in matrigel as described above using X-VIVO medium (#881024, Lonza) supplemented with 10% Fetal Bovine Serum (#10270106, Thermofisher), 1% Pen/Strep (#15140122, Thermofisher) and 1% Glutamax (#35050061, Thermofisher). MET induction was started once the cells were seeded into the matrigel. The culture time of the spheroids was of 9 days. On the 10th day, the co-culture was initiated by addition of the PBMCs. For this a cryostock of PBMCs was thawed into a cell-repelent well plate (#665970, Greiner) at a concentration of 1 *×* 10^6^ cells/mL. After an overnight recovery time, the PBMCs were stained with cell proliferation dye eFluor 670 or eFluor 450 (#65-0840-85,#65-0842-85, Thermofischer) according to manufacturer’s protocol before seeding. 1 *×* 10^5^ cells were added to each well of the matrigel spheroids culture. To activate T cells, Transact (#130-111-160, Miltenyi) was added to the culture medium in a 1:100 dilution. Cells were co-culture at 37 *^◦^*C and 5% CO_2_ e for 72 hours.

#### 3. Sample preparation for flow cytometry analysis

To get a homogeneous single cell suspension, clusters were dissociated by using a combination of mild mechanical disruption and enzymatic treatment with TrypLE (#12604013, Thermofischer). In brief, the cultures were washed with PBS and incubated with TrypLE for approximately 5 min at 37 *^◦^*C. The matrigel disc was dispersed with a 200*µ*L-pipette by pipetting up and down and doing circular movements in the well. The content of each well was transfered separately to one eppendorf where further TrypLE was added and the incubation was prolonged up to 30 min with intermediate checks in the microscope to observe the appearence of single cells. Afterwards, the samples were centrifuged and re-suspended directly in the staining buffer. Annexin-V and Propidium Iodide (PI) (#640914, Biolegend) staining was performed following the manufacturer’s protocol.

#### 4. Flow cytometric analysis of apoptotic cells

Stained cells were re-suspended in PBS and measured in a LSRFortessa™ FACS instrument (BD Biosciences). Flow cytometry data were analyzed using the FlowJo 10.10.0 software. To determine the apoptosis on a single cell level, single viable (PI negative) cells were analyzed for the expression of the apoptosis marker Annexin V. Among the Annexin V positive cells, the tumor cell population was identified due to being eFluor450/670 negative.

#### 5. Immunostaining of matrigel 3D cultures and imaging

Live imaging was done prior to fixation, by using a Zeiss confocal LSM980 airyscan 2 confocal microscope and a Plan-Apochromat 10x/0.45 air objective. Immediately afterwards, the co-culture was fixed with 100 *µ*L of 3.7% formaldehyde/PBS for 1 h in a rocking shaker at room temperature. Permeabilization was done afterwards with 0.5% Triton X-100/PBS during 45 min with constant rolling at room temperature. Blocking was performed overnight at 4 *^◦^*C with a blocking buffer of PBS and 1% BSA, 0.2% Triton X-100 and 0.05% Tween. After blocking, the primary antibody was incubated overnight at 4 *^◦^*C as well, diluted into blocking buffer. Three washings of 1 h each with PBS on constant rolling were performed in between the primary and the secondary antibody incubation. The secondary antibody incubation was done overnight, the solution was prepared as well in blocking buffer and DAPI (#28718-90-3, Sigma Aldrich) was added to the solution in the same concentration used for other stainings. After the DAPI/secondary antibody incubation was done, a similar washing step was performed again before the mounting of the sample into clearing solution^92^. Imaging was performed with a ND1 Nikkon spinning disk confocal microscope with a 25x/1.05 Silicon immersion Nikon objective from the Physics of Life microscopy facility.

#### 6. Image analysis

The number of apoptotic foci and PBMC-infiltration was quantified manually in each cell cluster. For this, clusters/spheroids of an average cross-sectional area of 5 *−* 7 *×* 10^4^ *µ*m^2^ were considered. Big clusters exceeding this size were divided into two independent clusters for the analysis to normalize the effect of spheroid size. Apoptotic foci were defined as regions of intense Cleaved Caspase-3 staining that were accompanied by a fragmented cell nuclei and absence of eFluor 670 signal, see Supplementary Fig. S6e. For PBMCs infiltration, the positive events were considered as those where the lymphocytes were found within the boundaries of a spheroid/cluster or directly bound to its surface, See Supplementary Fig. S6f.

### P. Statistical analysis

For statistical analysis of qPCR measurements, see Fig. 1b and Fig. 7b, five and four individual biological replicates were used respectively and three technical replicates were obtained for each. The value of ‘n’ refers to the number of biological replicates. The ΔCt values for the control condition were averaged across all replicates, and the resulting mean was used to calculate the ΔΔCt values for each replicate in the induced condition. The fold change for the control was determined by calculating the ΔΔCt for each replicate, relative to the mean ΔCt of the controls.

In assays with adherent cells, see Fig. 2, 3 and S3, the sampling was done randomly in different regions of the well. Each recorded frame constituted a data point in the statistics. For area constraint experiments, see Fig. 2h-j, each single cell was imaged separately and each cell constitutes a data point. In either case, ‘n’ refers to the total number of frames or cells sampled. These were sampled from at least two independent biological replicates.

Data on 3D cultures are presented in Fig. 4, S4, 5, S5, 7 and S7. There, either single spheroids were sampled (non-degradable gels) or low magnification confocal images of a large field of view were recorded (degradable gels) to capture cell nuclei in dense clusters as well as in sparse intermediate spaces (Fig. 4j and 7e). In either case, specimens were screened to provide a representative overview of the culture and the field of view was chosen at random in different regions of the well. In non-degradable gels each spheroid constitutes a data point. In degradable gels, each frame recorded was considered a data point. ‘n’ represents the total number of spheroids/frames sampled. These were obtained from at least two independent biological replicates with at least 2 gels per condition per experiment.

During image analysis of co-culture experiments, see Fig. 6 and S6, clusters were selected at random in different areas of the well to obtain a representative population. Each cluster constitutes a data point. The total number of clusters sampled, denoted as ‘n’, was obtained from three biological replicates, each containing at least two wells per condition. For flow cytometric analysis of apoptotic cells, see Fig. 6 and S6, at least three wells per conditions were used on each experiment. Each well was collected separately and constituted a technical replicate. The variable ‘n’ refers to the total number of technical replicates, or wells, obtained from a total of 3 independent biological replicates.

## ACKNOWLEDGMENTS

EFF acknowledges financial support from the Deutsche Forschungsgemeinschaft under Germany’s Excellence Strategy, EXC-2068-390729961, Cluster of Excellence Physics of Life of TU Dresden. Furthermore, EFF was supported by the Deutsche Forschungsgemeinschaft (DFG, German Research Foundation) by the grant FI 2260/7-1 and by the Heisenberg program – project number 495224622 (FI 2260/8-1). The authors extend their gratitude to Carsten Werner, along with Alison Le and Nelly Rein from his team, for supplying the materials used in the PEG-Heparin hydrogels. In addition, the authors thank the PoL Light Microscopy Facility for excellent support. Further, they thank Jan Kuhlmann for the provision of the ovarian cancer cell line ES-2. GD thanks Sanika Jahagirdar for providing the protocols for immunostaining of matrigel-grown tumor spheroids.

## Notes

https://gitlab.com/polffgroup/met_analysis

### Competing Interest Statement

The authors have declared no competing interest.

