## Supplementary information for "Mesenchymal-epithelial transition reduces proliferation but increases immune evasion in tumor spheroids"

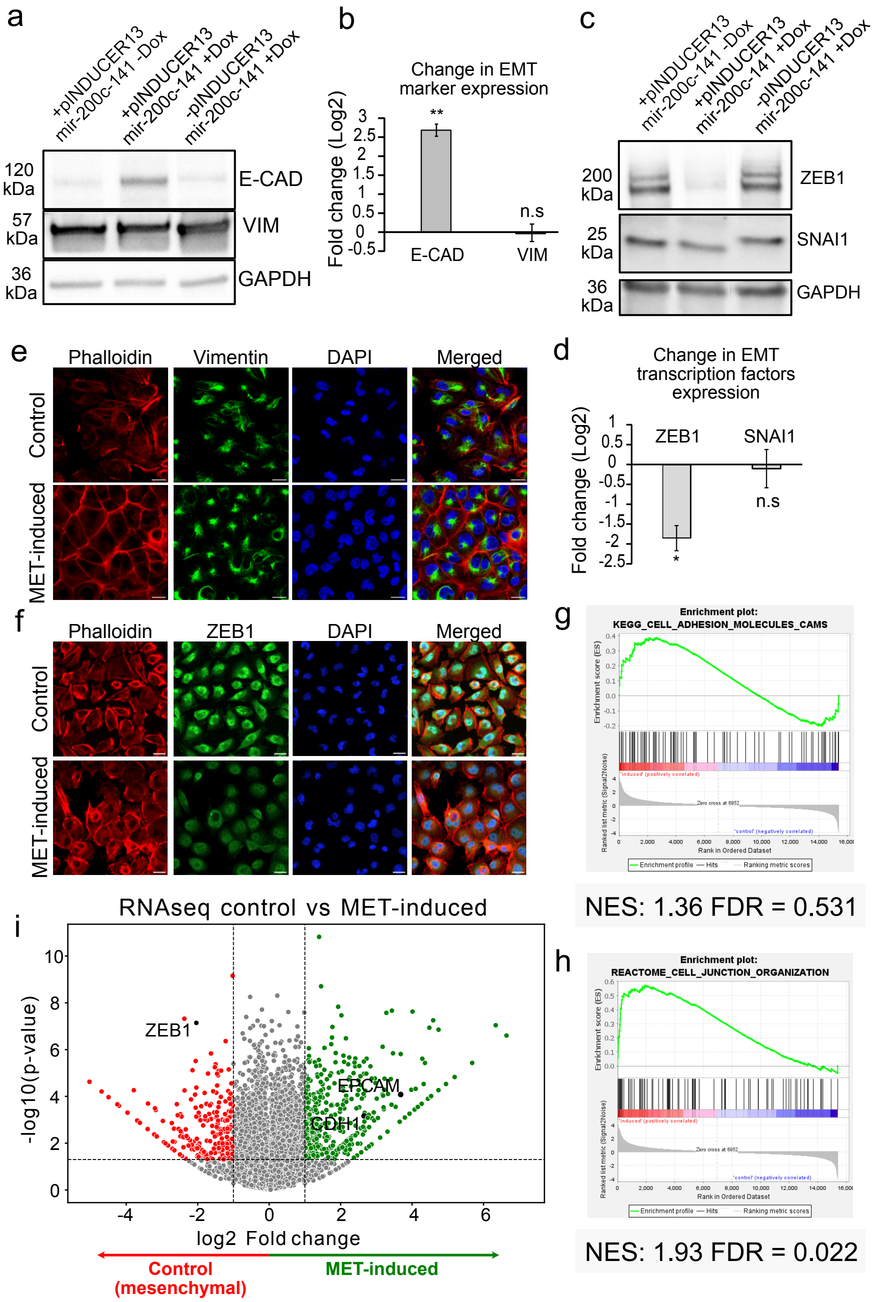


FIG. S1. Overexpression of the miR-200c-141 microRNA cluster leads to MET in mesenchymal breast cancer cells MDA-MB-231. a, b) Western-blot analysis of EMT markers E-cadherin and Vimentin without and with MET induction in transfected cells (first two columns) and in doxycyclin-treated untransfected cells (third column). Data were obtained from three independent biological replicates. Significance was tested with a two-tailed unpaired T-Tests against a Fold change of 0. c, d) Western-blot analysis of EMT-associated transcription factors SNAI1 and ZEB1 without and with MET induction in transfected cells (first two columns) and in doxycyclin-treated untransfected cells (third column). Data were obtained from three independent biological replicates. Significance was tested with a two-tailed unpaired T-Test against a fold change of 0. e, f) Immunofluorescence for mesenchymal marker Vimentin (e) and transcription factor ZEB1 (f) in control cells (top row of each panel) and induced cells after 4 days of induction (bottom row of each panel). Scale bar = 20 $\mu$m. g, h) GSEA analysis for gene sets shows that the induced phenotype is in line with MET. We tested enrichment for the gene sets 'reactome_cell_junction_organization' (reactome.org, R-HSA-446728, doi: 10.3180REACT_20676.1) (g) and 'KEGG_adhesion_molecules_CAMS' (KEGG PATHWAY: hsa04514) (h) from the canonical pathways of the 'C2: curated gene sets' collection. NES = Normalized Enrichment Score, FDR = False Discovery Rates. i) Volcano plot showing statistically significant changes in total gene expression upon overexpression of miR-200c-141. Red data points correspond to genes with reduced expression while green data points represent genes with increased expression.

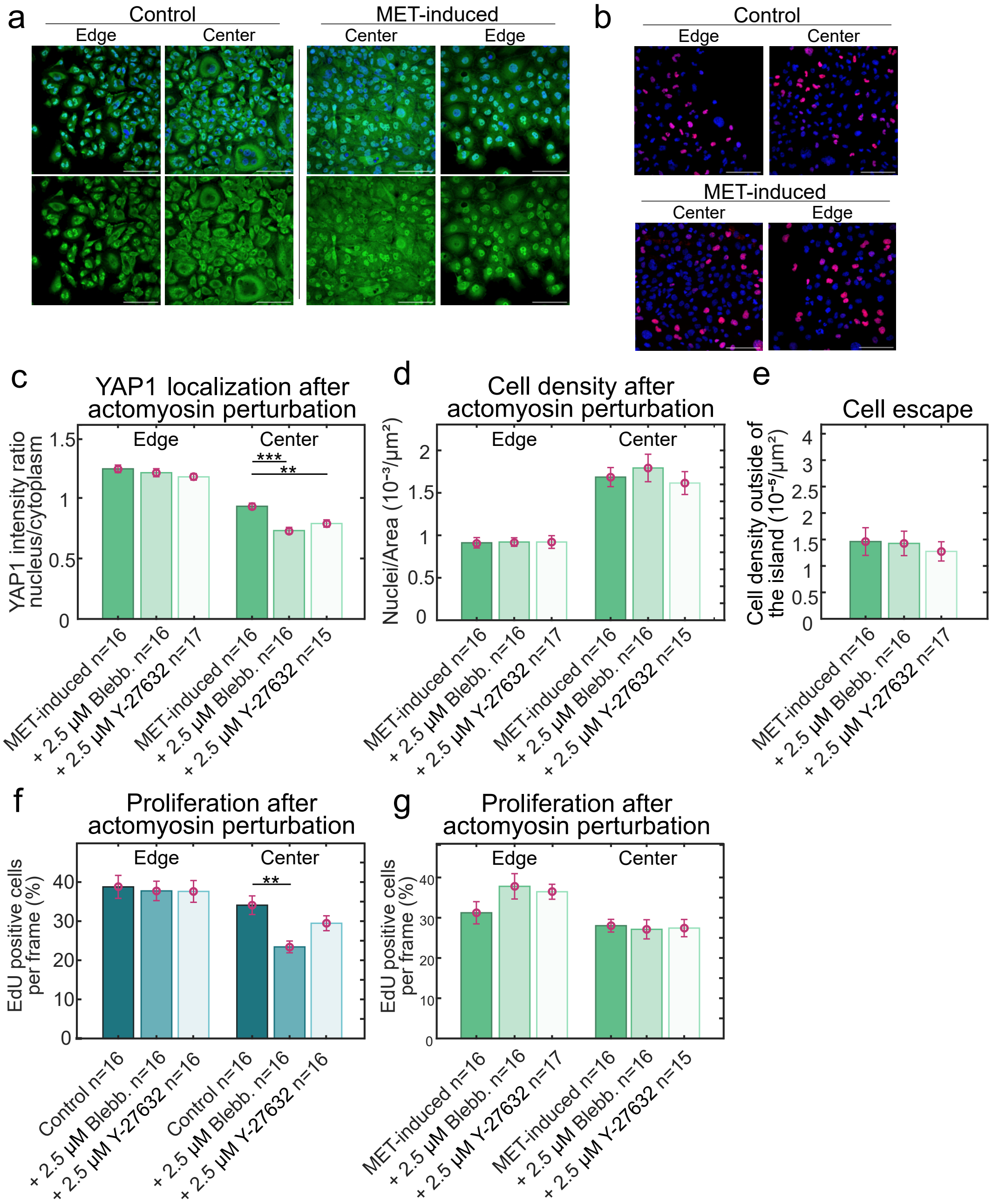


FIG. S3. MET-induction reduces proliferative signaling in self-organized dense islands of MDA-MB-231 cells with open boundaries. a, b) Close-ups of MDA-MB-231 cells in control (a = left panel, b= top panel) and MET-induced (a = right panel, b= bottom panel) conditions acquired at the center or edge of self-organized cell islands. These are example images of the frames used for the quantification of both YAP1 nuclear accumulation and proliferation rate. a) YAP1 immunofluorescence (green). Top row: DAPI staining of nuclei and YAP1 immunofluorescence. Bottom row: only YAP1. Scale bar = 50 $\mu$m. b) EdU fluorescence (red) and DAPI staining of cell nuclei in regions at the edge and the center of cell islands. Nuclei of cells that underwent S-phase during the incubation time incorporated EdU and correspondingly appear red in the images. Scale bar = 50 $\mu$m. c,d) YAP1 nucleus-to-cytoplasm ratio (c) and cell density (d) both in the center and edge regions of self-assembled cell islands formed by MET-induced MDA-MB-231 with and without actomyosin inhibitors treatment. e) Cell nuclei density outside of cell islands. Each data point corresponds to cell counts in a peripheral region of on average 1 mm$^2$. f) Percentage of EdU positive cells in islands of MDA-MB-231 cells (f: control, g: MET-induced) with and without indicated actin-cytoskeletal perturbations in both center and edge regions of the islands. For c, d, e and f, each data point corresponds to one frame recorded. ‘n’ represents the total number of frames recorded, these were obtained from 2 independent biological replicates. Bar heights show the mean and error bars the standard error of the mean. Significance was tested using Mann-Whitney-U-Test and $p \leq 0.05$.

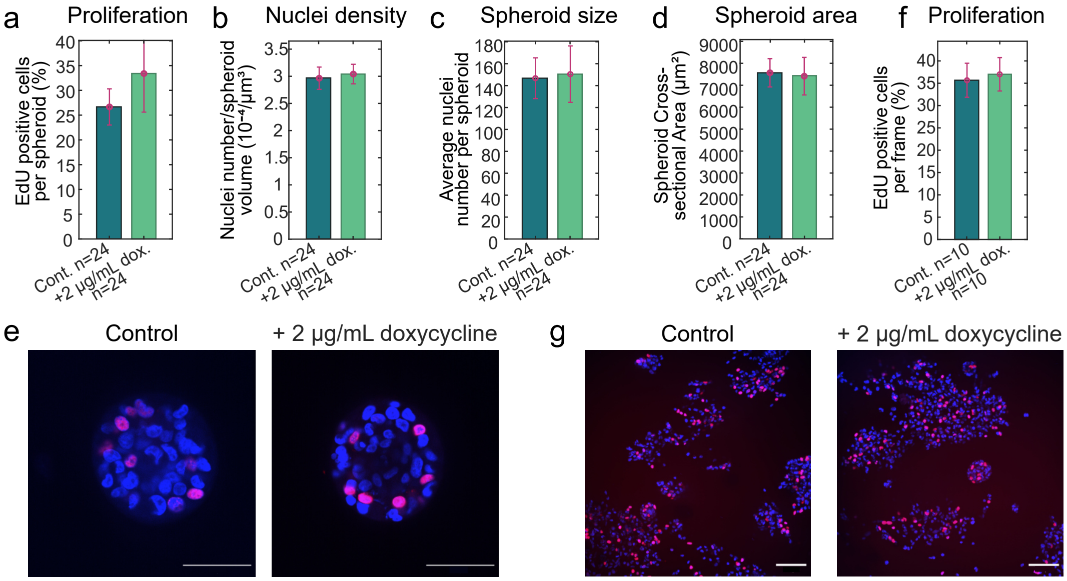


FIG. S4. Doxycycline does not change proliferation of untransfected MDA-MB-231 in 3D culture. a-d) Quantification of the percentage of EdU positive nuclei (a), nuclei density (b), spheroid sizes, in terms of nuclei numbers (c) and spheroid cross-sectional area (d) in non-degradable PEG-Heparin hydrogel spheroids of untransfected MDA-MB-231 with and without doxycyline in the medium. The nuclei density was calculated in terms of number of nuclei per spheroid volume assuming a sphere-shaped spheroid. f) Quantification of the percentage of EdU positive cells in 3D cultures of untransfected MDA-MB-231 in degradable PEG-heparin hydrogels. e,g) Representative confocal images of untransfected MDA-MB-231 cells in (e) non-degradable PEG-Heparin hydrogels or (g) degradable PEG-heparin hydrogels in control condition (left) or with doxycyline in the medium (right). Nuclei appear blue due to DAPI staining and EdU positive cells are red. e) Scale bar = 50 $\mu$m. g) Scale bar = 100 $\mu$m. For panels a-d, each data point corresponds to one spheroid, 'n' represents the total number of spheroids sampled. In f, each data point corresponds to a recorded frame. For panels a-f, bars heights show the mean and error bars the standard error of the mean. Significance was tested using Mann-Whitney-U-Test and $p \leq 0.05$.

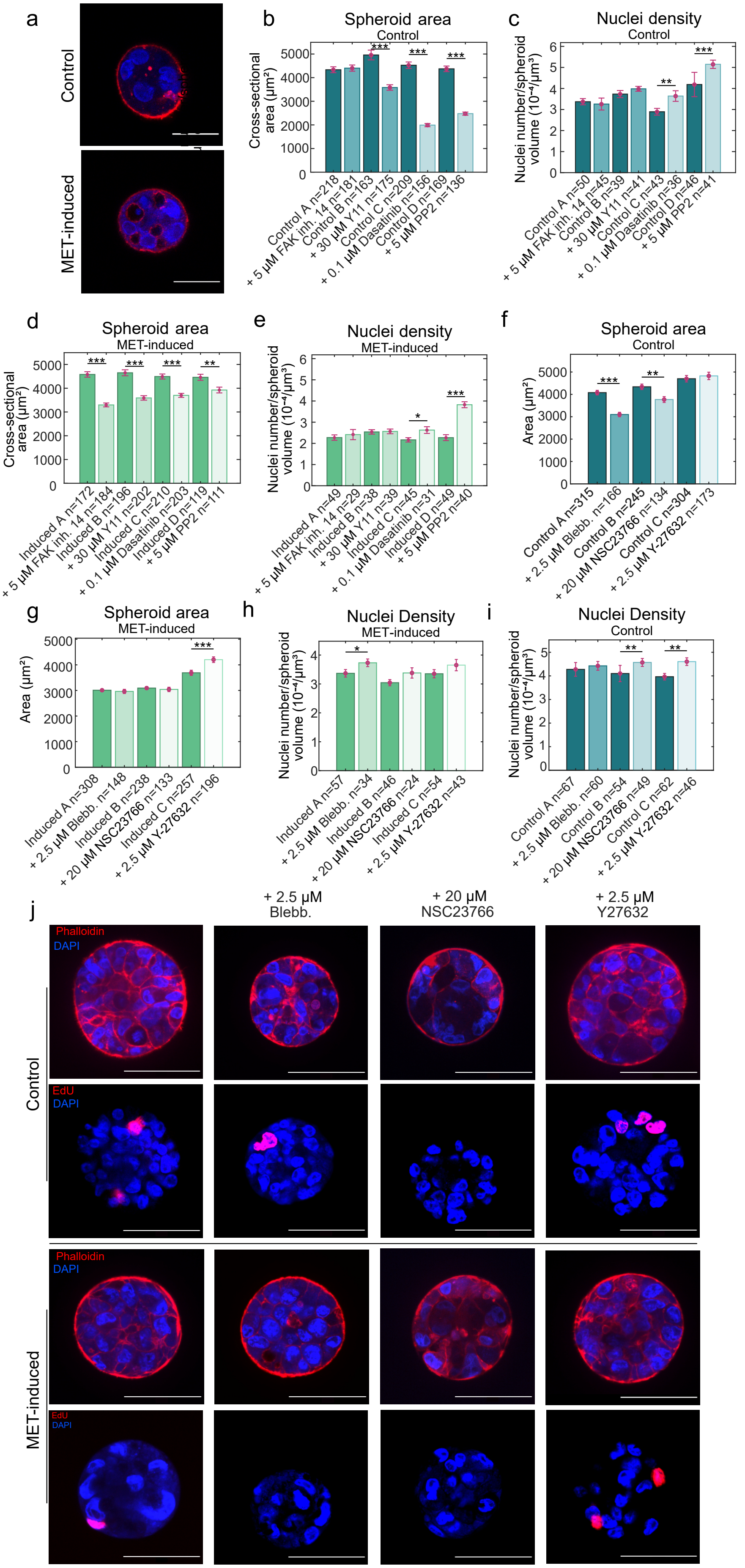
a
Control
MET-induced
b
Spheroid area
Control
Cross-sectional area (μm²)
Control A n=218
+ 5 μM FAK inh. 14 n=181
Control B n=163
+ 30 μM Y11 n=175
Control C n=209
+ 0.1 μM Dasatinib n=156
Control D n=169
+ 5 μM PP2 n=136
c
Nuclei density
Control
Nuclei number/spheroid volume (10⁻⁴/μm³)
Control A n=50
+ 5 μM FAK inh. 14 n=45
Control B n=39
+ 30 μM Y11 n=41
Control C n=43
+ 0.1 μM Dasatinib n=36
Control D n=46
+ 5 μM PP2 n=41
d
Spheroid area
MET-induced
Cross-sectional area (μm²)
Induced A n=172
+ 5 μM FAK inh. 14 n=184
Induced B n=196
+ 30 μM Y11 n=202
Induced C n=210
+ 0.1 μM Dasatinib n=203
Induced D n=119
+ 5 μM PP2 n=111
e
Nuclei density
MET-induced
Nuclei number/spheroid volume (10⁻⁴/μm³)
Induced A n=49
+ 5 μM FAK inh. 14 n=29
Induced B n=38
+ 30 μM Y11 n=39
Induced C n=45
+ 0.1 μM Dasatinib n=31
Induced D n=49
+ 5 μM PP2 n=40
f
Spheroid area
Control
Area (μm²)
Control A n=315
+ 2.5 μM Blebb. n=166
Control B n=245
+ 20 μM NSC23766 n=134
Control C n=304
+ 2.5 μM Y-27632 n=173
g
Spheroid area
MET-induced
Area (μm²)
Induced A n=308
+ 2.5 μM Blebb. n=148
Induced B n=238
+ 20 μM NSC23766 n=133
Induced C n=257
+ 2.5 μM Y-27632 n=196
h
Nuclei Density
MET-induced
Nuclei number/spheroid volume (10⁻⁴/μm³)
Induced A n=57
+ 2.5 μM Blebb. n=34
Induced B n=46
+ 20 μM NSC23766 n=24
Induced C n=54
+ 2.5 μM Y-27632 n=43
i
Nuclei Density
Control
Nuclei number/spheroid volume (10⁻⁴/μm³)
Control A n=67
+ 2.5 μM Blebb. n=60
Control B n=54
+ 20 μM NSC23766 n=49
Control C n=62
+ 2.5 μM Y-27632 n=46
j
+ 2.5 μM Blebb.
+ 20 μM NSC23766
+ 2.5 μM Y27632
Phalloidin
DAPI
EdU
Control
MET-induced

FIG. S5. Perturbations of focal adhesion-associated kinases and actomyosin affect proliferation differently on mesenchymal and MET-induced MDA-MB-231. a) Representative images of control (top) and MET-induced (bottom) MDA-MB-231 spheroids after two weeks of culture in non-degradable PEG-Heparin hydrogels in the absence of the gel-adhesion mediator RGD. Scale bar = 20 $\mu$m. b-e) Changes of tumor spheroid growth upon treatment with FAK and Src inhibitors. Growth was quantified according to cross-sectional area (b, d) and nuclei density (c, e) in control (b, c) and MET-induced (d, e) conditions. For area measurements, the maximal cross-sectional area was chosen from confocal z-stacks. Nuclei density was calculated in terms of number of nuclei per spheroid volume assuming a sphere-shaped spheroid. f-i) Changes of tumor spheroid growth upon treatment with actomyosin inhibitors. Growth was quantified according to cross-sectional area (f,g) and nuclei density (h,i) in tumor spheroids of control (f, i) and MET-induced MDA-MB-231 (g,h). Cytoskeletal perturbations of actomyosin were applied through treatment with: 2.5 $\mu$M Blebbistatin, 20 $\mu$M NSC 23766 RAC1 inhibitor, or 2.5 $\mu$M Y-27632 Rho kinase inhibitor. For panels b-i, every tumor spheroid analyzed gave rise to one data point. 'n' represents the total number of spheroids sampled, these were obtained from two (b-e) or three (f-i) independent biological replicates. Bar heights show the mean and error bars the standard error of the mean. Significance was tested using Mann-Whitney-U-Test and $p \leq 0.05$. j) Representative confocal images showing tumor spheroids of control MDA-MB-231 and MET-induced MDA-MB-231 with and without cytoskeletal perturbation of actomyosin. Top row: F-actin looks red by phalloidin staining and nuclei are stained by DAPI (blue). Bottom row: Edu-positive nuclei appear read indicating that the cells went into S-phase during the incubation period. Scale bar = 50 $\mu$m.

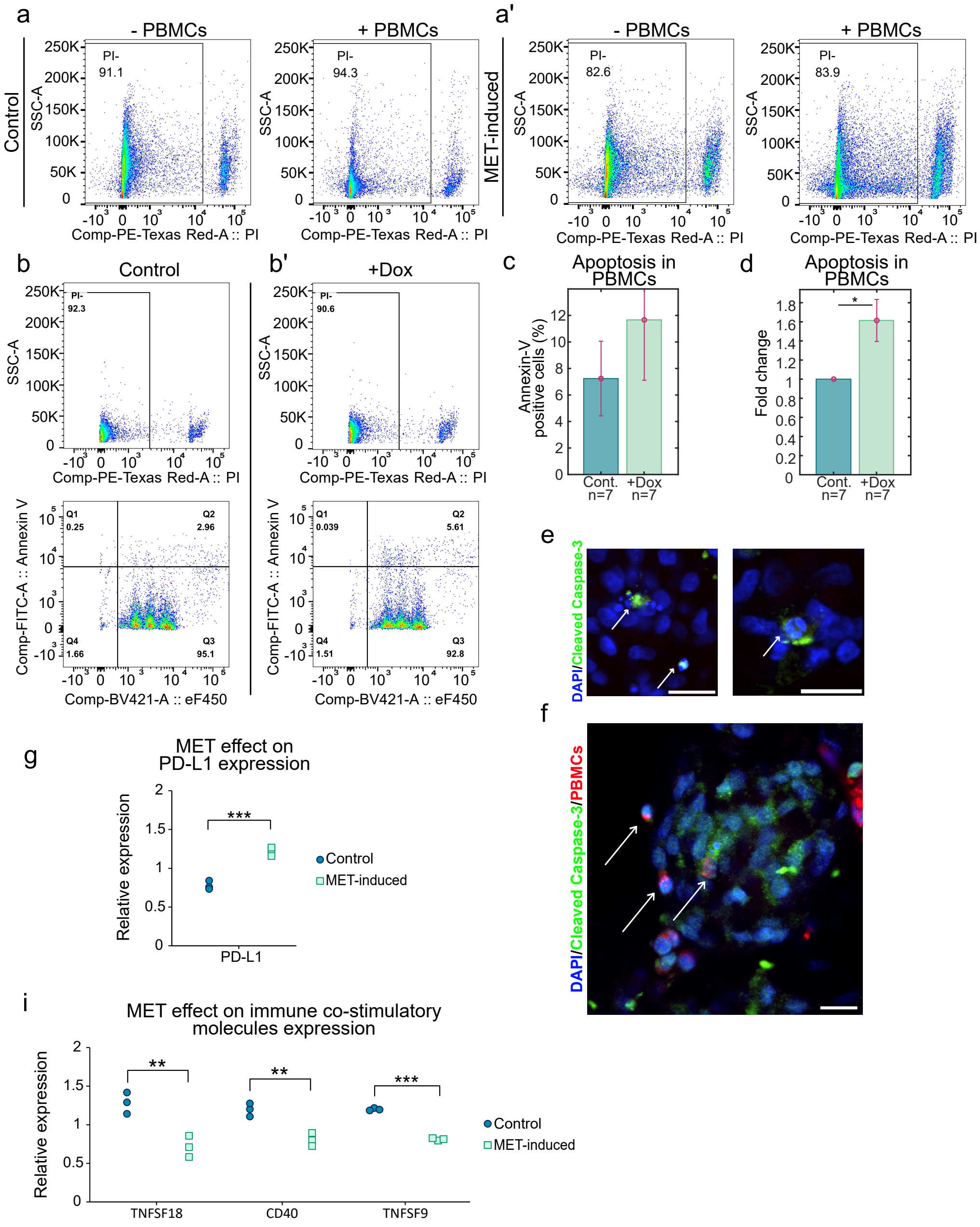

a
- PBMCs
+ PBMCs
Control
SSC-A
PI-
91.1
94.3
Comp-PE-Texas Red-A :: PI
a'
MET-induced
82.6
83.9
b
Control
92.3
b'
+Dox
90.6
Comp-FITC-A :: Annexin V
Comp-BV421-A :: eF450
Q1
0.25
Q2
2.96
Q3
95.1
Q4
1.66
0.039
5.61
92.8
1.51
c
Apoptosis in PBMCs
Annexin-V positive cells (%)
Cont. n=7
+Dox n=7
d
Fold change
e
DAPI/Cleaved Caspase-3
f
DAPI/Cleaved Caspase-3/PBMCs
g
MET effect on PD-L1 expression
Relative expression
PD-L1
Control
MET-induced
i
MET effect on immune co-stimulatory molecules expression
TNFSF18
CD40
TNFSF9

FIG. S6. MET enhances survival rate of tumor spheroids in the presence of peripheral blood mononuclear cells. a,a') Flow cytometry plots showing propidium iodide (PI) fluorescence versus side scatter (SSC-A) for cell suspensions containing MDA-MB-231 (extracted from dissociated tumor spheroid cultures) with or without additional PBMCs. Black frames demonstrate gating for cell viability which was done before analysis for apoptosis by Annexin-V staining. b,b') Flow cytometry plots with PBMCs from cultures without cancer cells in control medium (b) or in the presence of 2 $\mu$g/mL doxycycline (b'). The top row shows side scatter plots versus PI fluorescence intensity and the gating strategy for cell viability (black box). In the bottom row, Annexin-V intensity versus eFluor 450 intensity is plotted. To obtain quantifications of apoptotic PBMCs the ratio of cell numbers in the two right quadrants (Q2 and Q3) was calculated. c) Percentage of Annexin-V positive PBMCs from the flow cytometry plots in panel b. Each well of culture constitutes a data point. 'n' represents the total number of technical replicates, these were obtained from 2 independent biological replicates. d) Fold change increase of apoptosis upon doxycycline addition to the medium. To calculate the fold change, the wells were paired according to their position in the well-plate. e) Confocal close-ups of tumor spheroids fixed and stained with DAPI (blue) and antibodies for Cleaved Caspase-3 (green). The latter was used for the quantification of apoptotic foci. The arrows point to examples that were considered as apoptotic foci: a bright green fluorescent signal in combination with a fragmented cell nucleus. Scale bar = 20 $\mu$m. f) Confocal close-ups of tumor spheroids fixed and stained with DAPI (blue) and antibodies for Cleaved Caspase-3 (green), and PBMCs labeled with cell proliferation dye eFluor670 (red), used for the quantification of PBMC infiltration. The arrows point to examples that were considered positive cases of infiltration: PBMCs (red dots) inside the tumor spheroid mass or close to its surface. Scale bar = 20 $\mu$m. g) RNAseq quantification of PD-L1 expression on MDA-MB-231 cells before and after MET induction in adherent cultures. Significance was tested using a two-tailed Student T-test. i) RNAseq quantification of the expression of immune co-stimulatory proteins TNFSF18, CD40, TNFSF9 on MDA-MB-231 cells before and after MET induction in adherent cultures. Significance was tested using a two-tailed Student T-test.

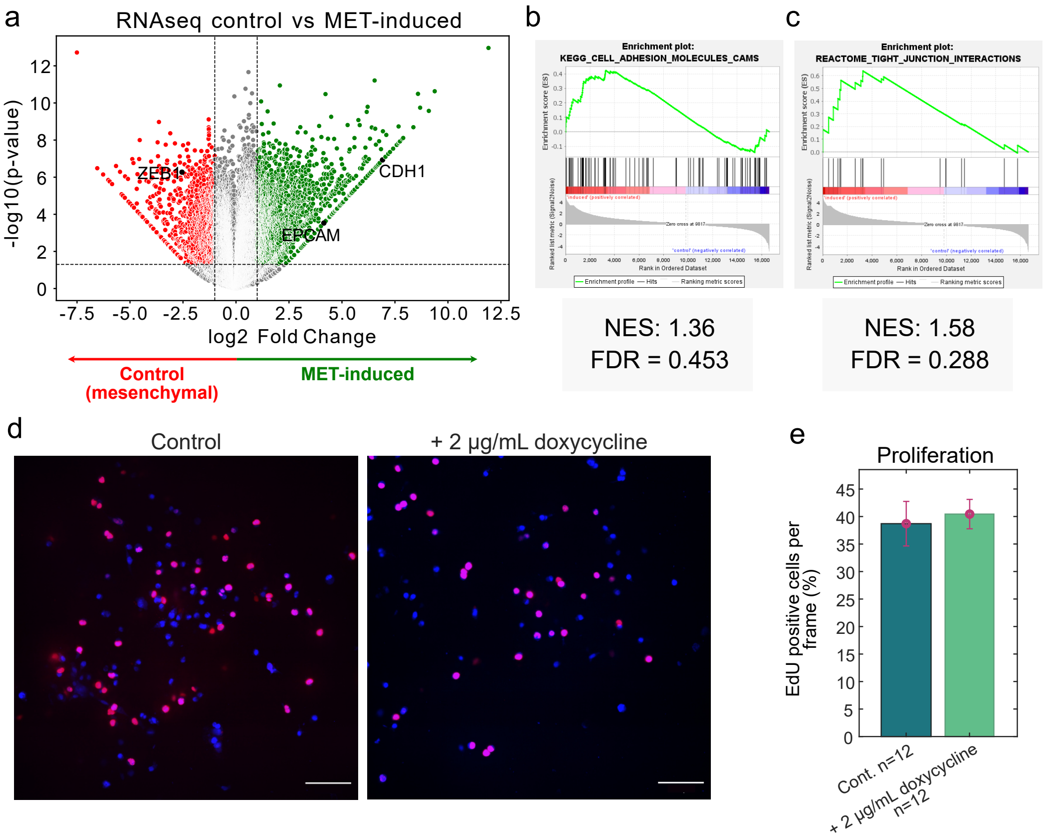


FIG. S7. Doxycycline does not change proliferation of untransfected ES-2 in 3D culture. a) Volcano plot showing statistically significant changes in total gene expression upon overexpression of miR-200c-141. Red data points correspond to genes with reduced expression while green data points represent genes with increased expression. b,c) GSEA analysis for gene sets shows that the induced phenotype is in line with MET. We tested enrichment for the gene sets 'KEGG_adhesion_molecules_CAMS' (KEGG PATHWAY: hsa04514) (b) from the canonical pathways of the 'C2: curated gene sets' collection and 'reactome_tight_junction_interaction' (reactome.org, R-HSA-420029) (c). NES = Normalized Enrichment Score, FDR = False Discovery Rates. d) Representative images of untransfected ES-2 cells grown in degradable PEG-Heparin hydrogels in control medium (left image) and with the addition of 2 $\mu$g/mL doxycyline (right image). Cell nuclei were stained with DAPI (blue) and the proliferation marker EdU (red). Scale bar =100 $\mu$m. e) Quantification of the percentage of EdU positive cells. 'n' represents the total number of frames sampled, these were obtained from 4 separate gels and 2 independent biological replicates. Bars correspond to mean and error bars are standard error of the mean. Significance was tested using Mann-Whitney-U-Test and $p \leq 0.05$.

| Component | Individual amount (µL) |
|---|---|
| 10X buffer | 1.5 |
| primer RNU6B | 2.39 |
| primer miR-200c | 2.39 |
| primer miR-141 | 2.39 |
| RNAse Inhibitor | 0.19 |
| Reverse Transcriptase | 1 |
| dNTPs | 0.15 |

TABLE S3. cDNA synthesis reactions components.

| Step | Temperature | Time |
|---|---|---|
| Reverse transcription | 16 °C<br>42 °C | 30 min<br>30 min |
| Stop reaction | 85 °C | 5 min |
| Hold | 4 °C | Hold |

TABLE S4. Cycling steps used in the cDNA synthesis reaction

| miR-141 | miR-200c | RNU6B |
|---|---|---|
| 14 µL of 20x primer for miR-141<br>140 µL TaqMan Master mix<br>107.38 µL H2O DNAse free | 14 µL of 20x primer for miR-200c<br>140 µL TaqMan Master mix<br>107.38 µL H2O DNAse free | 14 µL of 20x primer for RNU6B<br>140 µL TaqMan Master mix<br>107.38 µL H2O DNAse free |

TABLE S5. Quantities of reactants used for quantitative PCR reactions for the detections of each microRNA.

| STEP | Temperature | Time | Cycles |
|---|---|---|---|
| UNG activation | 50 °C | 2 minutes | 1 |
| Enzyme activation | 95 °C | 10 minutes | 1 |
| Denature<br>Anneal/extend | 95 °C<br>60 °C | 15 seconds<br>60 seconds | 40 |

TABLE S6. Cycling steps done in the quantitative PCR reactions.
